# Zinc- and redox-dependent allosteric regulation of the *Helicobacter pylori* CemR transcription factor

**DOI:** 10.64898/2026.11.26.769429

**Authors:** Jakub Muraszko, Piotr Szczepanowski, Mateusz Noszka, Katarzyna Kapczyńska, Krzysztof Skowronek, Aleksander Czogalla, Michal Tracz, Artur Krezel, Anna Zawilak-Pawlik

## Abstract

The human gastric pathogen *Helicobacter pylori* senses redox conditions *via* a pleiotropic atypical response regulator, CemR (HP1021). The protein’s activity is regulated by the redox state of cysteine residues and Zn^2+^ (i.e., Zn-Cys redox switch signaling), but the molecular mechanisms underlying ROS detection and signal transduction remain uncharacterized. Here, we demonstrate that CemR comprises three domains: an N-terminal receiver (RD), a central domain (CD), and a C-terminal DNA-binding domain (DBD). Cysteine residues of the RD and CD/DBD domains sense redox conditions via 2-Cys and 1-Cys mechanisms, respectively, and are required for Zn²⁺ coordination, thereby affecting DNA-binding activity. Yet each domain influences the protein’s activity in its own characteristic way. Under reducing conditions, CemR binds DNA most effectively, and reduced cysteine residues in the CD/DBD domains are crucial for this interaction. Zn²⁺ binding primarily impacts the RD domain, inhibiting DNA binding when CemR is reduced. This suggests that structural changes in the RD domain transmit signals to the CD/DBD domain, regulating transcriptional activity. RD domain cysteine residues are also required for reversible redox regulation of CD/DBD activity. Because the predicted dimer structure shows no direct intra-interactions between protomer domains, allosteric regulation likely controls CemR activity in response to redox and zinc levels, possibly via the signaling α-helix and the unstructured regions linking RD to CD and CD to DBD, respectively. CemR is unique in that it evolved from a two-component system, but its receiver-like domain uses cysteine-dependent redox chemistry and metal coordination, rather than phosphorylation-dependent signaling.

**Bullet points:**

- CemR consists of N-terminal RD, central CD, and C-terminal DBD domains, each controlled by redox.
- Allosteric interactions between RD and CD/DBD are critical for CemR activity control.
- Reduction of the RD’s disulfide bond increases CemR affinity for Zn²⁺, reducing affinity for DNA.
- Oxidation of Cys residues in the CD/DBD domains reduces CemR’s affinity for DNA.
- Reversible redox control of CD/DBD activity depends on Cys residues in RD.

## 1. Introduction

Bacteria are constantly exposed to oxidative stress from reactive oxygen species (ROS) generated endogenously as by-products of aerobic metabolism or from exogenous sources. ROS are highly toxic to cells because they eventually damage biomolecules. To manage the response to oxidative stress, bacteria use redox switch regulators, i.e., proteins that sense redox imbalance within the cell and undergo structural changes that modify their activities [1,2]. This enables the cell to transduce signals and ultimately initiate a response to redox stress.

Redox switches are typically standalone proteins [1,2]; however, some evolved from two-component systems (TCS) response regulators [3]. Redox switches, which are often transcription regulators, sense ROS via redox modification at the amino acid residue (e.g., hydrogen peroxide-inducible gene activator, OxyR, modified at cysteine residues [4] ) or by the metal cofactor (e.g., Fe-S cluster in fumarate and nitrate reduction regulator, FNR [5,6]) or both, the metal cofactor and the amino acid residue can be engaged in ROS sensing (e.g., peroxide sensing regulator, PerR, oxidized at the histidine residue via cofactor Fe^2+^-dependent Fenton reaction [7]). Cysteine residues, specifically their sulfhydryl (-SH, thiol) groups, are predominantly employed by redox switches as a sensory mechanism for rapid activity regulation [8–10]. Thiol oxidation leads to sulphenic acid formation (-SOH), which is unstable and may further react with low-molecular-weight compounds, such as glutathione (S-glutathionylation) in a 1-Cys mechanism, or another cysteine residue, forming a disulfide S-S bond in a 2-Cys mechanism. Increased thiol oxidation leads to sulfinic (-SO_2_H) and sulfonic (-SO_3_H) acids, with the latter being irreversible. Redox switches often use zinc ions to co-regulate their activity. Under reducing conditions, Zn^2+^ can coordinate cysteine residues, stabilizing thiolate ions and lowering the p*K*_a_ of the cysteine residue. Under oxidative conditions, thiolate ions are oxidized, releasing Zn^2+^. Cysteine residues then form a disulfide bond, altering the structure and modulating protein activity (e.g., DNA affinity). At the same time, the released Zn^2+^ may also propagate the redox signal within the cell as a second messenger of the redox conditions. It should be noted that, beyond redox signaling, cysteine residues, whether complexed with Zn^2+^ or not, also perform other versatile roles in proteins, including enzymatic and structure-stabilizing functions [11,12].

*Helicobacter pylori* is a gram-negative, microaerobic, obligate human pathogen [13]. It inhabits the stomach’s harsh environment, being the leading cause of peptic ulcers and gastric cancer, and the second leading cause of death from cancer worldwide [13]. The *H. pylori* genome (1.67 Mbs, 1576 ORFs in the 26695 strain) encodes only three sigma factors (a housekeeping σ^80^, and alternative σ^54^ and σ^28^)[14] and approximately 30 signal transduction factors, which form a regulatory, often overlapping network that helps *H. pylori* maintain homeostasis and respond to environmental signals such as acidic pH, nutrient and metal availability, or response to immune cells[14–18]. *H. pylori* employs post-transcriptional regulation to control gene expression, since approximately 900 non-coding RNAs were identified in *H. pylori* 26695 [19,20]. Significant overlap between transcriptional and post-transcriptional gene expression control, and a poorly characterized landscape of post-translational modifications [21–24], lead to a poor understanding of many *H. pylori* processes, including oxidative stress response.

*H. pylori* is exposed to oxidative stress during transmission and infection [25–27]. On the one hand, the pathogen uses a limited repertoire of enzymes or compounds to combat ROS and lacks the general stress-response regulator RpoS and glutathione–glutaredoxin (GSH/Grx) reduction system (reviewed in [28]). In contrast, *H. pylori* employs non-classical strategies to either combat or benefit from oxidative stress. It can take up the antioxidant ergothioneine (thiourea derivative of histidine), which, based on studies in mice, may facilitate *H. pylori* infection [29,30]. As shown recently, *H. pylori* senses and migrates towards strongly bactericidal HClO^-^ - rich niches seeking inflamed tissues – an efficient source of nutrients, especially iron [31,32]. ROS-induced DNA mutations may increase *H. pylori* genetic variation and adaptability to human hosts, especially during the acute infection phase [33,34]. Under oxidative stress, the pathogen actively takes up large amounts of foreign DNA via a ComB type IV secretion system, possibly using it as a “shield” to protect its own DNA from oxidative damage [35], which promotes chronic infection [36]. Moreover, H. pylori incorporates foreign DNA into its genome to evolve during human infection [37]. Thus, the bacteria seem to benefit from oxidative stress by effectively managing its stress response.

We recently demonstrated that an atypical, orphan HP1021 response regulator, renamed Campylobacteria energy and metabolism regulator (CemR) [38], functions as a redox-switch protein [1,39]. CemR, one of the least-characterized H. pylori transcriptional regulators, is not essential for H. pylori viability, but its knock-out reduces H. pylori growth rate [39,40] (see [41] for a comprehensive review of H. pylori transcriptional regulation). CemR acts as both an activator and a repressor, controlling gene expression under microaerobic conditions and in response to oxidative stress triggered by aerobic conditions [39,42,43].

The molecular mechanism underlying CemR activity control remains enigmatic. A previously published model indicated its two-domain structure [39]. The N-terminal domain is similar to the phosphoacceptor receiver REC domain of the two-component system (TCS) response regulators, but lacks the conserved aspartic acid residue that receives a phosphate group from the histidine kinase (HK); such domains are called pseudo-receiver domains, or more specifically, aspartate-less receiver (ALR) domains [44–47]. The C-terminal DNA-binding domain comprises a helix-turn-helix motif (HTH) [39]. *H. pylori* CemR protein contains six cysteine residues at positions C27, C51, and C56 in the N-terminal domain, and C216, C233, and C270 in the C-terminal domain; only C27 is conserved in *Campylobacterial* species beyond *H. pylori*. Cysteine residues in CemR mediate redox-dependent modulation of protein structure, thereby influencing CemR’s DNA- and Zn^2+^-binding activities. Specifically, their oxidation and Zn^2+^ binding reduce the affinity of CemR for DNA [39]. Previous studies have shown that CemR coordinates Zn^2+^ in a 1:1 stoichiometry, with the affinity ranging from *K*_d_ 10^-9^-10^-8^ M for reduced CemR to 10^-7^-10^-6^ for oxidized CemR [39], which is higher than the affinity of other typical Zn^2+^-controlled regulators, such as zinc uptake regulator, Zur (*K*_d_ of 10^-15^)[48].

In this study, we demonstrate that the CemR protein comprises three domains - the N-terminal receiver RD, the central CD, and the C-terminal DNA-binding DBD - with the central domain being distinct, both sequentially and structurally, from previously characterized protein domains. The cysteine residues in the RD and CD/DBD domains sense redox conditions and coordinate Zn²⁺, thereby influencing CemR DNA-binding activity, with each domain affecting the protein’s activity in its own characteristic way. In detail, CemR binds DNA with the highest affinity under reducing conditions, with the reduced cysteine residues in the CD/DBD domain being most critical for CemR’s efficient interactions with DNA. However, CD/DBD cysteine residues are not directly involved in DNA interactions, but they are likely important for maintaining CemR’s proper, redox-responsive structure. CD/DBD cysteine oxidation is reversible; however, it depends on the RD domain cysteine residues. Zn^2+^ exerts most pronounced regulatory effect through the RD domain, inhibiting CemR-DNA binding when CemR is reduced. Therefore, redox-induced structural changes in the RD domain in the presence of Zn^2+^ are transmitted to the CD/DBD domain, thereby regulating CemR transcriptional activity. The predicted dimer structure shows no interactions between the RD and CD/DBD domains; thus, we propose that allosteric mechanisms regulate CemR activity in response to redox conditions and possibly zinc levels. Although detailed structural changes remain to be revealed, the signaling α-helix and the unstructured regions linking RD to CD and CD to DBD are likely involved in CemR allosteric regulation.

## 2. Materials and methods

### 2.1. Materials and culture conditions

The strains, plasmids, and proteins used in this work are listed in Supplementary Table S1. The primers used in this study are listed in Supplementary Table S2. *H. pylori* was cultivated at 37 °C under microaerobic conditions (5% O_2_, 10% CO_2,_ and 85% N_2_) generated by the jar evacuation-replacement method (PetriSphere Anaerobic System). *H. pylori* plate cultures were grown on Columbia blood agar base medium supplemented with 7% defibrinated horse blood (CBA-B). The liquid cultures were prepared in Brucella Broth (BB) (Becton Dickinson), each containing 10% fetal bovine serum (FBS) (Biowest) (BB-FBS); the cultures were shaken at 140 rpm. All *H. pylori* cultures were supplemented with an antibiotic mix [49]. If necessary for selecting *H. pylori* mutants, appropriate antibiotics were used with final concentrations (i) kanamycin 15 µg/ml, (ii) chloramphenicol 8 µg/ml. *Escherichia coli* DH5α and BL21 were used for cloning and recombinant protein synthesis, respectively. The *E. coli* MC1061 strain was used to propagate the plasmid for transformation into *H. pylori*. If necessary for selecting *E. coli,* appropriate antibiotics were used with final concentrations (i) kanamycin 50 µg/ml and (ii) ampicillin 100 µg/ml.

### 2.2. RNA isolation

*H. pylori* cultures (12 ml BB-FBS) of N6 wild-type and mutant strains were grown under microaerobic conditions to an OD_600_ of 0.8-1.0. Immediately after opening the jar, 1 ml of the non-stressed culture was added to 1 ml of RNAprotect Bacteria Reagent (Qiagen), vortexed, and incubated for 5 min at room temperature. In parallel, the cultures were moved to aerobic conditions for 25 min (air atmosphere, 37 °C, 140 rpm orbital shaking). After oxidative stress, samples were collected as described for non-stressed cells. After 5 min incubation with RNAprotect Bacteria Reagent, collect bacteria by centrifugation (7,000 × g, 5 min, RT). RNA was isolated immediately after bacterial collection, or the bacterial pellet was stored at −80 °C for no longer than three months. RNA was isolated using the GeneJET RNA Purification Kit (Thermo Fisher Scientific; K0731) according to the manufacturer’s protocol and treated with RNase-free DNase I (Thermo Fisher Scientific). Next, the RNA was further purified using the GeneJET RNA Purification Kit to remove any remaining DNase I. A NanoDrop Lite spectrophotometer and agarose gel electrophoresis were used to determine the quality and quantity of RNA. The isolated RNA was stored at -80 °C for up to one month.

### 2.3. Quantitative polymerase chain reaction

Reverse transcription was performed using 500 ng of RNA in a 10 µl reaction mixture with the iScript cDNA Synthesis Kit (Bio-Rad). 2.5 µl of 1:10-diluted cDNA was added to 7.5 µl of SsoAdvanced Universal SYBR Green Supermix (Bio-Rad) and 400 nM each of the forward and reverse primers, for a final reaction volume of 15 µl. The RT-qPCR program was performed as previously described [39] . The following primer pairs were used: P1-P2, *gluP*; P3-P4 *fecA3* (Table S2 and [39]). The relative mRNA abundance for each gene was determined by comparing mRNA levels to those of *H. pylori* 16S rRNA (P5-P6 primer pair). The RT-qPCR was performed using RNA isolated from four independent *H. pylori* cultures to generate a gene transcription dataset.

### 2.4. Protein expression and purification

The purification of recombinant Strep-tagged CemR proteins was performed as described earlier [39], with one significant modification. Namely, buffer W (100 mM Tris–HCl, pH 8.0; 300 mM NaCl and 1 mM EDTA), used at purification steps and as a solution to prepare elution buffer E (buffer W with 5 mM desthiobiotin), was demetallized using Helex 100 resin (1422822, Bio-Rad) prior to use. The purified fractions were stored at -20 °C in buffer E diluted with glycerol to a final concentration of 50%.

### 2.5. Electrophoretic mobility shift assay (EMSA)

EMSA was performed as described previously [39]. The *oriC*2 region was amplified by PCR using P7-P8 primer pairs and a specific pori2_1 template (Supplementary Tables S1 and S2), yielding the FAM-*oriC*2 probe (317 bp, 35% GC). The control nonspecific DNA probe (212 bp, 35% GC, a fragment of the HP0180 *H. pylori* 26695 gene, not bound by CemR [39]), was amplified by PCR in two steps. In the first step, an unlabeled DNA fragment was amplified using primer pair P9–P10. Primer P9 contained overhangs complementary to the P11 Cy5-labeled primer. The unlabeled fragments were purified and used as templates in the second round of PCR using the Cy5-labeled P11 primer and the unlabeled P10 primer used in the first step. CemR protein samples (80 mM) prepared in EMSA buffer (50 mM Tris–HCl, pH 8.0; 100 mM NaCl and 0.2% Triton X-100) were incubated at 37 °C with sequential addition of reagents, each for 10 min. The following compounds were added: 5 mM Tris(2-carboxyethyl)phosphine (TCEP, LS-3405.0025, Iris), 0.39 µM Zn^2+,^ and 1 mM ethylenediaminetetraacetic disodium salt (EDTA). Subsequently, FAM-labeled DNA (5 nM) was added, and samples were incubated for an additional 30 min before analysis. To assess the reversibility of diamide-induced oxidation of the CemR protein, protein samples (40-160 mM) in EMSA buffer were first incubated with FAM-labeled DNA (5 nM) for 10 min at 37 °C. The samples were then treated with 1 mM diamide and/or 5 mM Tris(2-carboxyethyl)phosphine (TCEP), with each treatment step lasting 10 min. The complexes were separated by electrophoresis on a 4% polyacrylamide gel in 0.5 × TB (1 × TB: 89 mM Tris, 89 mM borate) at 10 V/cm in the cold room (approx. 10 °C). The gels were analyzed by a Typhoon 9500 FLA Imager and ImageQuant software.

### 2.6. Surface plasmon resonance (SPR)

The Biacore system (Biacore T200, GE Healthcare) was used to study DNA-protein and protein–protein interactions by SPR [50]. SPR response values are expressed in resonance units (RU), and for most molecules, 1 RU corresponds to binding 1 pg of the molecule per mm^2^ of the chip surface. The analysis was as described before [39], with minor modifications. Biotinylated *oriC2* probe (346 bp, 35 %GC) was obtained by PCR with P12-P13 primer pair using the pori2_1 plasmid template (Supplementary Tables S1 and S2). The nonspecific DNA probe (206 bp, 35% GC) was amplified by PCR in two steps. In the first step, an unlabeled DNA fragment was amplified using primer pair P29–P30 and *Campylobacter jejuni* 11168 genomic DNA as a template. Primer P29 contained overhangs complementary to the P31 biotin-labeled primer. The unlabeled fragments were purified and used as templates in the second round of PCR using the biotin-labeled P31 primer and the unlabeled P30 primer used in the first step. (Supplementary Tables S1 and S2). DNA was immobilized on a Series S Sensor Chip SA (approx. 100 RU) according to the supplier’s instructions (GE Healthcare). The measurements were performed in Tris buffer (50 mM Tris–HCl, pH 8.0; 100 mM NaCl; and 0.2% Tween 20) at a continuous flow rate (15 μl*/*min). When indicated, CemR protein samples were reduced with 5 mM DTT. At the end of each cycle (180 s association phase followed by 180 s dissociation), bound proteins were removed by washing with 0.05% SDS (30 s), and the flow channels were equilibrated with Tris buffer until the baseline stabilized. The BIA evaluation 4.1 and Excel 2019 software programs were used for data analysis.

### 2.7. Monitoring HP1021 thiol redox state in vitro

To monitor the redox state of CemR *in vitro*, a differential thiol trapping analysis was performed using the thiol-reactive probe 4-acetamido-4’-maleimidylstilbene-2,2’-disulfonic acid (AMS, A485, Thermo Fisher Scientific) and iodoacetamide (IA, 16125, Merck). CemR (40 μl, 170 μM in buffer W (100 mM Tris-HCl pH 8.0, 300 mM NaCl)) was precipitated with an equal volume of 20 % trichloroacetic acid (TCA), washed with 100% ice-cold acetone, and dried. CemR was subsequently suspended in 40 μl of labeling buffer (300 mM Tris-HCl pH 7.5, 2% SDS). When indicated, the labeling buffer contained 20 mM TCEP, 2 mM tetramethylazodicarboxamide (diamide, D3648, Merck), 20 mM IA, 2 mM AMS, 6 M guanidinium hydrochloride (GuHCl), or 10 mM hydrogen peroxide (1.07209.0250, Merck). If multiple chemicals were used to treat CemR in subsequent reaction steps, each step was quenched by TCA precipitation, as described above, before proceeding to the next redox/labeling step. All reactions were carried out for 30 min. at 37 °C and were protected from light. The samples were supplemented with Laemmli buffer without any reducing agent, heated at 95 °C, and separated by electrophoresis on a 10% SDS-PAGE gel. The resolved gels were visualized by TCE-UV staining [51].

### 2.8. SDS-PAGE and Western blotting

Proteins were separated by electrophoresis on SDS-PAGE gels and visualized by the TCE-UV method [51] using a Gel Doc XR+ (Bio-Rad). The gels were transferred to nitrocellulose membranes and then incubated with an anti-HP1021 antibody [52] and anti-rabbit IgG (rabbit IgG HRP-linked whole Ab, NA934, GE Healthcare). The membranes were visualized by chemiluminescence, and images were captured using a ChemiDoc (Bio-Rad).

### 2.9. Ellman’s assay

Quantification of cysteine residues in a reduced thiol state in CemR was performed by colorimetric reaction of sulfhydryl groups with 5,5⍰-dithio-bis-(2-nitrobenzoic acid) (DTNB, Ellman Reagent, D8130 Merck). Analysis was performed according to the protocol for the Ellman Reagent (22582, Thermo Scientific). A standard curve was generated as a function of cysteine hydrochloride monohydrate concentration (0.25 mM to 1.5 mM), freshly prepared by serial dilution in the reaction buffer (0.1 M Na_3_PO_4_, pH 8.0, 1 mM EDTA). Reactions contained 625 μl of reaction buffer, 12.5 μl of DTNB (4 mg/ml), and 62.5 μl of cysteine standard or protein sample of the known concentration (C_CemR_, 10-180 mM). Samples were mixed and incubated at room temperature for 15 minutes. 200 μl of the reaction solution was added to a 96-well plate in triplicate. Absorbance was measured at 412 nm with a CLARIOstar (BMG LABTECH). The molar concentration of sulfhydryl groups (C_SH_) was calculated from the standard curve, and then the number of sulfhydryl groups per protein molecule (N_SH_) was calculated by dividing C_SH_ by C_CemR_.

### 2.10. Protein thermal stability

The CemR protein thermal unfolding analysis was performed by the Prometheus NT.48/Panta (NanoTemper), which uses nano-Differential Scanning Fluorimetry (nanoDSF)[53,54]. The thermal unfolding profiles of proteins were analyzed by monitoring changes in tryptophan fluorescence as proteins unfolded. The purified recombinant proteins (15 μM in 45 μl of Tris buffer (50 mM Tris-HCl pH 8, 300 mM NaCl)) were incubated in 1.5 ml reaction tubes for 30 minutes at 30 °C. When indicated, the protein samples contained TCEP (5 mM). 10 μl of sample was loaded into a capillary (Prometheus Standard Capillaries, NanoTemper), and then inserted in duplicate to the Prometheus NT.48/Panta. Thermal unfolding experiments were carried out in the range of temperature range from 25 °C to 95 °C, with a heating rate of 0.5 °C per minute. Data were analyzed using Panta Control 1.4 software, and inflection points indicating the melting temperature of the analyzed proteins were calculated. The analysis was repeated at least two times.

### 2.11. Zincon assay

To estimate the Zn^2+^-binding properties of CemR, a competition assay with 2-carboxy-2⍰-hydroxy-5⍰-sulfoformazyl-benzene monosodium salt (Zincon) was performed similarly as described [39]. Stock solutions of Zincon (96440, Merck) were prepared in DMSO (9.25 mg/ml). The affinity of Zincon for Zn^2+^ was analyzed by titrating 100 μM Zincon in buffer Z (100 mM Tris-HCl, pH 8.0; 300 mM NaCl; 5 mM desthiobiotin; and 10 μM ZnSO_4_, with 5 mM TCEP present when indicated) with CemR diluted in buffer E (see CemR purification protocol), in a 1-cm path length quartz cuvette and reading the absorbance at 618 nm at 25°C using a Hitachi U-2900 spectrometer. The absorption of the Zn^2+^-Zincon complex was transformed to the species concentration using its molar absorption coefficient, 225,970 M^-1^ cm^-1^. The dissociation constant of the Zn^2+^-Zincon complex at pH 8.0 is 2.14 × 10^-7^ M, as previously determined[55].

### 2.12. Bottom-up mass spectrometry

CemR wild-type protein (100 µg) was precipitated with 10% TCA. The precipitate was dissolved in 100 μl of Tris-urea buffer with NEM (8 M urea, 100 mM Tris-HCl, pH 6.5, 10 mM N-ethylmaleimide (NEM, Merck E3876)) and incubated for 1 hour at 37 °C in the dark. The protein was applied to a 10 kDa MWCO filter (VWR 516-0229) equilibrated twice with 200 μl 50.6 mM ammonium bicarbonate buffer (AmBic) and washed twice with 200 μl AmBic buffer. The CemR protein was digested overnight on-filter with trypsin at 37 °C (0.5 µg trypsin/100 µg CemR). The digested peptides were eluted from the filter by centrifugation into Eppendorf tubes (10,000 × *g*, 10 min, 4 °C). One hundred microliters of 0.1% formic acid (FA) were applied to the top of the filter and centrifuged into the peptide-containing Eppendorf collection tube (10,000 × *g*, 10 min, 4 °C). The peptide-containing filtrate was lyophilized and used for LC-ESI-MS analysis. Before analysis, the sample was reconstituted in an aqueous solution containing 0.1% FA and 3% acetonitrile (ACN) at 5 pmoles/µl of protein.

LC-ESI-MS was performed on an M-Class Acquity nanoUPLC linked to a Synapt XS HDMS (Waters). Mobile phase A consisted of H_2_O + 0.1% FA, while mobile phase B consisted of ACN + 0.1% FA (LC-MS grade solvents from WITKO). Two independent CemR preparations were investigated, with 2.5 pmoles of the protein sample injected each time. An 8-40% B 40 min linear gradient at a 300 nl/min flow rate was applied for sample separation on a C18 75 μm x 100 mm analytical column kept at 45 °C with a 3-minute sample trapping step before analytical column separation. MS data were collected in ion mobility DDA (HD-DDA) at an MS survey scan rate of 600 ms and an MS/MS scan rate of 200 ms, both in a 50–2,000 m/z range. Analyzer Mode was set to Resolution (∼45,000 FWHM at 785.84 m/z). Top 6 precursors with a charge of 2+, 3+ or 4+ were selected for MS/MS fragmentation with an m/z-specific CE ramp (LMS = 20 V, LME = 27 V, HMS = 52 V, HME = 62 V). MS/MS time was limited to 200ms. MS/MS data stripping based on the precursor’s drift was enabled. Source conditions were fine-tuned and kept constant. A leucine-enkephalin solution was acquired as a lock mass, and correction was applied post-acquisition.

Raw files were processed using PLGS v3.0.3 (Waters), and the deconvoluted data were exported as an .mgf query list, and searched using Mascot Server v2.8 (Matrix Science). To avoid combinatorial explosion, an iterative searching sequence was designed in Mascot Daemon. Search #1 was performed against an Escherichia coli reference proteome databank (UP000000625) to which CemR, trypsin, and human keratin sequences were appended. N-ethylmaleimide (C) and Oxidation (M) were set as variable modifications for this search. Search #2 was performed solely against the CemR sequence with N-ethylmaleimide (C) and Oxidation (M) as variable modifications, and an intact disulfide bridge (Xlink: disulfide[I] (C)) as a variable crosslinker. Search #3 used the same databank as Search #1, but had Oxidation (M), S-sulfenylation (C), S-sulfinylation (C), and S-sulfonylation (C) set as variable modifications. For each search, the peptide and fragment mass error tolerances were set to 15 ppm and 25 ppm, respectively, and two trypsin miscleavages were allowed. A PSM match significance threshold (peptide expected value) of 0.05 was used, with only unmatched queries being passed for a subsequent search. CemR peptides from Mascot .csv output files containing peptide-level match information from each search were combined into a separate Microsoft Excel sheet (Supplementary file 3) and inspected manually.

### 2.13. Intact and Top-down MS

Recombinant CemR samples were acidified to 0.1% FA and centrifuged (10,000 × *g*, 5 min). The supernatant was collected into a fresh vial and placed on the system. LC-ESI-MS was carried out on an M-Class Acquity UPLC system coupled to a Synapt XS HDMS (Waters). Mobile phase A consisted of H_2_O + 0.1% FA, 0.05% TFA, while mobile phase B consisted of ACN + 0.1% FA, 0.05% TFA. 2.8 pmol of protein was injected, desalted online, and a four-minute 10–80% B linear gradient was applied for sample separation on an Acquity UPLC BEH C4 300 Å, 1.7 μm, 1 mm x 50 mm column, which was kept at 80 °C. For intact protein mass, MS1 data were collected at a scan rate of 1s through a 500-2000 m/z range in positive polarity and TOF resolution mode. The spectra of the protein peak were combined, background-subtracted, and deconvoluted using MaxEnt1. For top-down MS, ion-mobility MS/MS data were collected for the 833.45 m/z ion of CemR (z = 44+) at a scan rate of 600 ms over a 300–2000 m/z range in positive polarity and TOF resolution mode. Spectra of the obtained mobility peak were split into 6 intervals of 10 bins each, and each interval was deconvoluted separately using MaxEnt3. A background threshold was applied to the resulting spectra (10% relative to BPI), and masses were combined into a single peak list. Ion-to-sequence matching was performed using ProSite Lite v1.4[56] at a 25 ppm threshold for BY and CZ ions. The disulfide modification between C51 and C56 was included in calculations. A match with a P-score of 2.3E−16 explaining 40% of fragments (by intensity) with a fragment mass error of 16.7 (ppm RMS) to the CemR sequence was obtained. All raw data were processed using the MassLynx V4.2 software. For every analysis, a Glufibrinopeptide B solution was acquired in the reference function, and the correction was applied during acquisition.

### 2.14. Statistics, reproducibility and data visualization

Statistical analysis was performed using Microsoft Excel or GraphPad Prism (version 8). The number of replicates and the statistical tests used for the analyses are specified in the figures’ captions. Unless otherwise stated, data were visualized by Microsoft Excel, GraphPad Prism, Corel package, and BioRender (https://www.biorender.com/).

### 2.15. Ethical issues

The HP1021 antibody used in Western blotting was raised in rabbits under the approval of the First Local Committee for Experiments with the Use of Laboratory Animals, Wroclaw, Poland (consent number 51/2012).

## 3. Results

### 3.1. CemR three-domain dimeric structure

Structure prediction analysis of *H. pylori* CemR in a 1:1 complex with Zn^2+^ performed by AlphaFold3 [57] indicated that the protein is actually composed of three domains and can form a dimer, as is typical for response regulators (Fig. 1A); the structure of a dimer is similar with and without Zn^2+^ (Fig. S1A). The N-terminal ALR-type domain (RD, 1-110 aa) is connected to the central domain (CD, 143-234 aa) via an extended α4 helix. The C-terminal DNA-binding domain (DBD) contains an HTH motif (272-292 aa)[58]. The BLAST protein sequence analysis revealed that the RD domain is most homologous to the AtoC-type regulator (COG2204) (Fig. S2). What distinguishes this domain from typical REC domains is the aforementioned substitution of the phosphoacceptor aspartic acid residue for a serine residue (S47), but also differences in amino acid sequence in conserved regions of the protein responsible for signal allosteric propagation (e.g., the T/S residue position required for “Y-T coupling” is changed to alanine (A72)) and structural changes of the conserved REC α2 helix into an unstructured region located between β2 and β3 of RD (Fig. 1 and Figs S2-S3)[46,59]. The C-terminal DBD domain is most homologous to the RocR-type regulator (COG3829) and contains a Fis-type DNA-binding HTH motif (IPR002197). Although the CD domain was classified as a P-loop-containing nucleotide triphosphate hydrolase in the TED analysis[58], this domain lacks the characteristic motifs, Walker A (G-X-X-X-X-G-K-[TS], X - any amino acid) and Walker B (hhhh[D/E]; h-hydrophobic amino acid)[60] (Fig. S2). According to BLAST analysis, the CD domain sequence differs from those of known proteins beyond homologous *Campylobacterial* CemR proteins. Significant parts of the CD and DBD domains remain unstructured. CemR contains no typical Zn^2+^ coordination motifs. The AlfaFold3 model places Zn^2+^ near residues H18, H21, and D22. The C51 residue of the RD and cysteine residues of the CD and DBD domains (collectively termed CD/DBD) are located in the unstructured, possibly flexible regions of the protein. The AF3 model of CemR predicts a putative intramolecular disulfide bond between C51 and C56, located at η1-α2 (AF-O25665-F1-v6, Fig. 1).

**Fig. 1.**
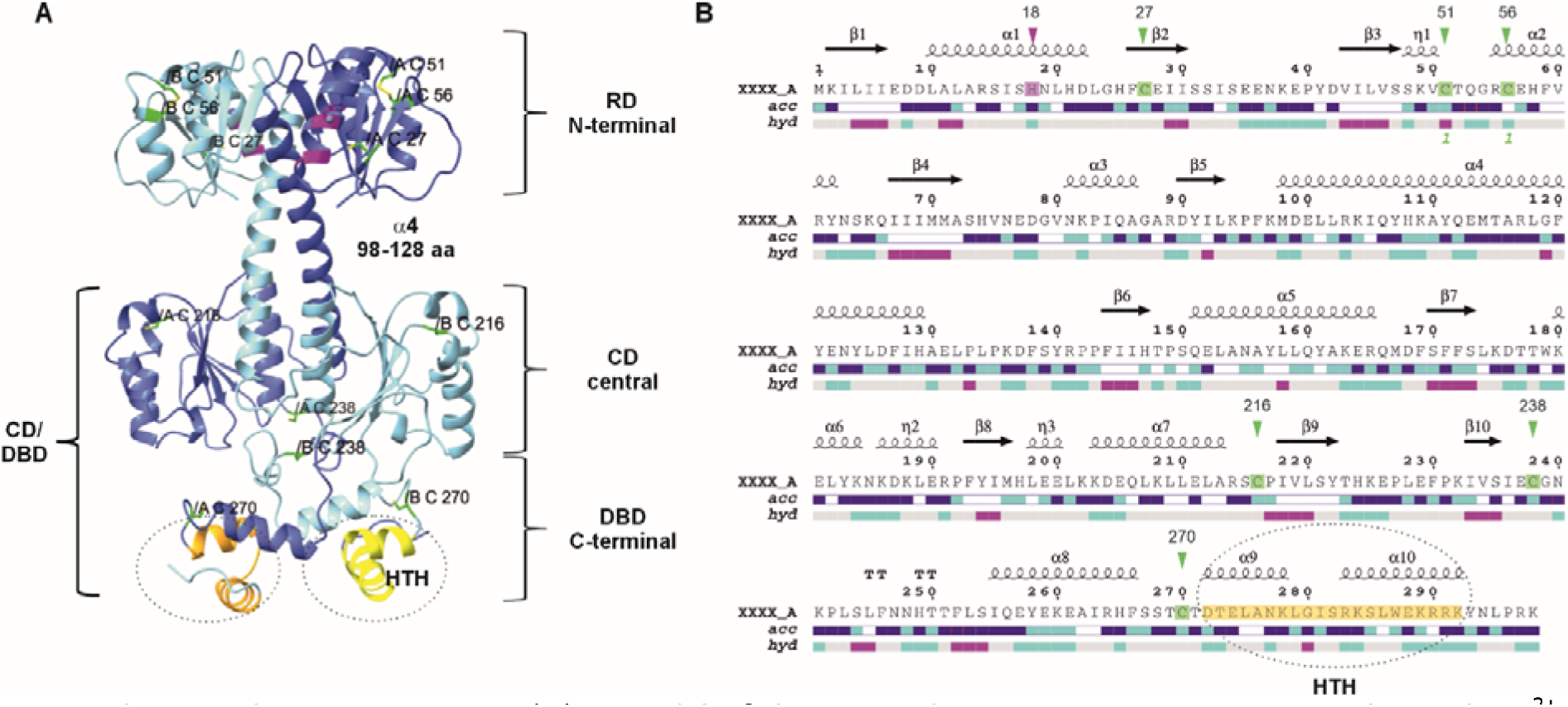
The *H. pylori* CemR protein. (**A**) A model of the CemR dimer structure in a 1:1 complex with Zn^2+^, predicted by AlphaFold3. Cysteine residues are highlighted in green, Zn^2+^ in grey, and proximal amino acids H18, H21, and D22 in magenta; the HTH motif is shown in orange and yellow (chains A and B, respectively). (**B**) CemR rendered by ENDstript2. Secondary structure elements are presented on top (squiggle, helices; arrows, strands; TT letters, turns; η, medium squiggle for an alpha helix); *acc*, solvent accessibility (blue, accessible; cyan, intermediate; white, buried); *hyd*, hydropathy (pink, hydrophobic; white, neutral; cyan, hydrophilic). Green „1" below the *hyd* bar indicates a disulfide bond predicted in CemR. Cysteine residues and His18, which is predicted to be involved in Zn^2+^ binding, are highlighted in green and magenta, respectively, and indicated by green and magenta arrowheads above the secondary structure elements. The HTH motif is highlighted in orange.

Size exclusion chromatography-multi-angle light scattering (SEC-MALS) analysis confirmed that CemR forms a stable dimer regardless of redox state (Fig. S1B). Thus, intermolecular disulfide bonding is not required for dimer formation and stability, consistent with a model in which the cysteine residues of the two protomers are not close to each other (Fig. 1).

### 3.2. RD and CD/DBD domains contain redox-reactive cysteine residues conforming 2-Cys and 1-Cys regulatory mechanisms

Because previous results indicated that the redox state of cysteine residues controls CemR activity, we purified a series of CemR muteins to identify the cysteine residues involved in redox control of CemR activity (Figs 2A and S4 and Table S1). Specifically, we replaced cysteine with alanine residues (C→A) in the RD or CD/DBD domains of muteins, designated RD-CA and CD/DBD-CA, respectively. Additionally, we created muteins lacking a conservative cysteine residue at position 27 (C27A) or the cysteine residues at positions 51 and 56 (C51A-C56A). A mutein in which all cysteine residues were replaced with alanine residues was previously constructed (C_less_, previously named ΔCys)[39]. We also constructed muteins representing only the N-terminus and the C-terminus (CemR_RD_, 1-127 aa and CemR_CD/DBD_, 119-298 aa, respectively). We Strep-tagged all proteins and purified them using the same protocol (Materials and Methods and Fig. S4). However, the Strep-tagged CemR_DBD_ variant was unstable, limiting its use in analyses (see below).

**Fig. 2.**
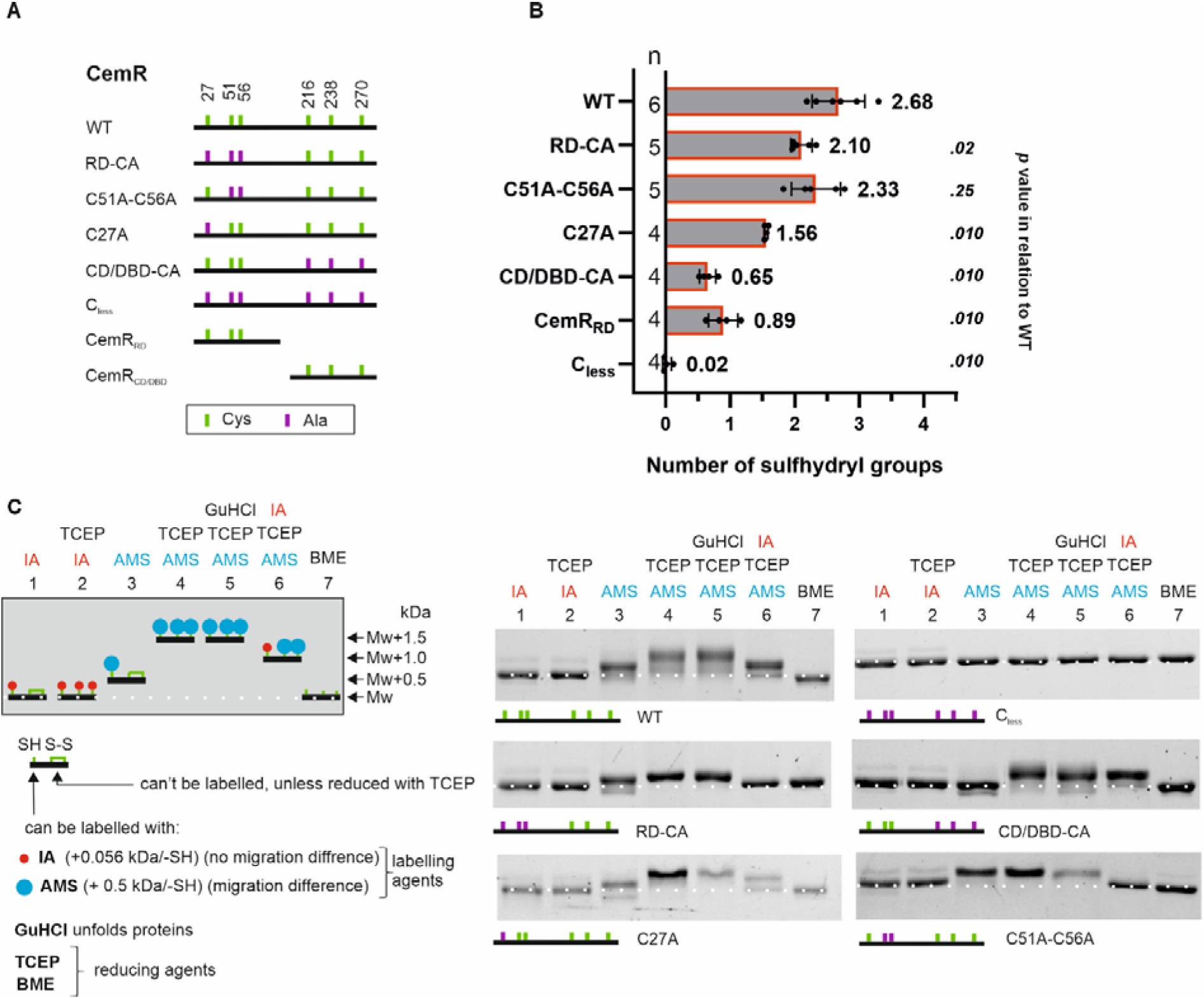
Analysis of CemR thiol redox state *in vitro*. (**A**) Schematic overview of the CemR variants used in this work. (**B**) Analysis of sulfhydryl groups in native, air-oxidized CemR muteins by labeling with Ellman reagent. Analysis was carried out in at least four independent replicates (n), with at least two technical replicates in each. Data are shown as mean ±SD from n replicates; n is specified on the left side of the bar for each protein variant. Mann-Whitney test was used to determine the *P* value. (**C**) Analysis of the CemR thiol redox state by differential thiol trapping. Left panel: schematic overview of the approach; right panel: the analysis of the thiol redox state in air-oxidized CemR. WT and its variants were incubated in Tris buffer containing reducing and/or labeling agents. GuHCl was added to the denture protein to expose the putative buried cysteine residues to reduction and labeling. The agents were added to the CemR protein sample in subsequent steps, as indicated above the gels (from top to bottom). The protein was precipitated with TCA after each step. CemR was resolved in a 10% SDS-PAGE gel under non-reducing conditions and visualized by the TCE method. Results for a broader range of molecular masses, including oxidized multimeric proteins, are presented in Figure S5. The white dotted line indicates the migration of unlabeled or IA-labeled CemR with no difference in migration. IA, iodoacetamide; AMS, 4-acetamido-4’-maleimidylstilbene-2,2’-disulfonic acid; TCEP, tris(2-carboxyethyl)phosphine; GuHCl, guanidine hydrochloride; BME, β-mercaptoethanol. Digital processing was applied equally across all images, including controls.

Next, we used Ellman’s reagent (5,5’-dithiobis-(2-nitrobenzoic) acid, DTNB) to detect sulfhydryl (-SH) groups present in the CemR wild-type and muteins and indirectly determine the number of cysteine residues oxidized by proteins’ exposure to air (air-oxidized proteins). The number of -SH groups in the wild-type CemR was determined to be 2.68±0.41, while no -SH group was detected in C_less_ protein (Fig. 2B). The number of -SH groups was higher in RD-CA (2.10 ± 0.16) than in CD/DBD-CA and CemR_RD_ variants (0.65 ± 0.13 and 0.89±0.23, respectively), indicating the presence of approx. one -SH group in the RD domain and two -SH groups in the CD/DBD domain. Exchanging C51 and C56 with alanine reduced CemR labeling by DTNB less than the C27A exchange (2.33 ± 38 and 1.56 ± 0.03 cysteine residues labeled, respectively), and the number of detected -SH groups in RD-CA and C51-C56 was similar to that in the WT protein. Thus, C51 and C56 are likely oxidized and, as predicted by AlphFold3, may form a disulfide bond, while C27 is likely reduced. Additionally, approximately one redox-reactive cysteine residue is located in the RD/DBD domain.

To analyze cysteine residue reactivity in more detail, differential thiol trapping was applied using iodoacetamide (IA) and 4\-acetamido-4\-maleimidylstilbene-2,2\-disulfonic acid (AMS) (Fig. 2C and Fig. S5). IA and AMS bind to -SH groups, increasing the molecular weight of a protein molecule by approximately 0.06 kDa and 0.5 kDa, respectively, for each labeled thiol group. Thus, IA blocks -SH groups without significant changes in molecular weight or migration on an SDS-PAGE gel, while AMS labels proteins and, due to the significant increase in molecular weight, retards their migration in a gel. Tris(2-carboxyethyl)phosphine (TCEP) was used to provide reducing conditions, while guanidinium chloride (GuHCl) was used to denature the protein’s structure, thereby uncovering potentially buried cysteine residues that were inaccessible for AMS. The CemR variants were labeled in a series of subsequent reactions, in a variable order depicted in Fig. 2C (see also Materials and Methods), and analyzed by SDS-PAGE. The C_less_ mutein was used as a control and, as expected, was not labeled by AMS under any conditions. All other air-oxidized CemR variants, except CD/DBD-CA, were efficiently labeled by AMS, indicating that these proteins contained -SH groups (Figs. 2C and S5B, lanes 2 vs 3). CD/DBD-CA was either unlabelled by AMS or, because of the small number of available -SH groups and the resulting small increase in molecular weight, its migration did not change significantly.

Nonetheless, the labeling analysis indicated that -SH groups are mainly located in the CD/DBD domain. The differences in migration between the AMS-labeled oxidized or reduced forms of protein variants were only observed for variants having C51 and C56 cysteine residues (WT, CD/DBD-CA, and C27A), indicating that these residues form the S-S bond, which is reduced by TCEP (Fig. 2C, lanes 3 *vs* 4 and 6). All -SH groups, either in the oxidized protein or after TCEP reduction, were likely surface-exposed, because GuHCl treatment did not reveal any additional -SH groups. Notably, diamide- or H₂O₂-oxidated recombinant proteins containing cysteine residues in the CD/DBD domains (WT, RD-CA) formed intermolecular disulfide bonds (CemR dimers), which did not occur in the CD/DBD-CA mutein lacking C-terminal cysteine residues (Fig. S5). The physiological significance of this oxidation is difficult to predict, as no disulfide-mediated dimer formation has been demonstrated *in vivo* under oxidative stress induced by anaerobic culture[39], and these dimers do not form *in vitro* in the protein isolated under non-reducing conditions. Yet, since cysteine residues are distantly located in the predicted structure, the results suggest structural mobility within these domains, consistent with the prediction of a high proportion of unstructured regions.

Finally, we performed a comprehensive mass-spectrometric analysis of two independent batches of the CemR wild-type protein. These included a derivatization-aided bottom-up analysis, as well as intact protein and top-down measurements without derivatization.

In the bottom-up analysis, we obtained nearly 80% protein sequence coverage and identified over 30 distinct CemR peptides. The obtained peptide maps covered 5 of the 6 cysteine residues present in the CemR sequence (C51, C56, C216, C238, and C270) (Fig. S6). The analysis yielded high-confidence spectral matches for a disulfide bond between C51 and C56, as well as for oxidation of C216 and C270 to sulfinic and sulfonic acids (-SO_2_H, -SO_3_H) (Fig. S7). Notably, despite the presence of feasible trypsin cleavage sites (R14 and K38, ) within the sequence, peptides corresponding to residues 14–38 were never identified. Therefore, no information on C27 could be gathered.

A major drawback of the bottom-up approach in protein analysis is the difficulty of translating peptide-level observations into accurate stoichiometric distributions of modification sites at the intact-protein level. Therefore, to assess the predominant proteoform of the recombinantly produced CemR, we measured its intact mass under denaturing conditions. The main peak (∼77% of TIC) corresponded to the mass of our CemR construct minus two hydrogen atoms (−5.5 ppm), suggesting a stable disulfide bond within the CemR monomer (Fig. S8). We ascribed the second-most-intense peak (∼8.3% of TIC) to in-source fragmentation, while the third (∼7.2% of TIC) comprised an unknown species, albeit unlikely to be oxygenation-related. Furthermore, the bottom-up analysis did not cover the C27 residue; however, given its high conservation, we sought to exclude its involvement in disulfide formation entirely. To do this, we tuned the CID of intact CemR (z = 44+) to obtain a fragment pattern in the vicinity of C27 and confirmed this residue as unmodified (Fig. S9).

Taken together, our mass spectrometry results provide strong complementary evidence that CemR protomers contain an intramolecular disulfide bond located in the RD domain between C51 and C56, while the remaining cysteine in this domain, C27, appears redox-unaffected. Furthermore, we showed that C216 and C270 in the CD/DBD domains may be susceptible to ROS modification. The CD/DBD ROS-modified residues are, however, undetectable at the intact-protein level, implying that these modifications are either artefactual or affect just a small pool of CemR molecules and occur only under specific conditions, as discussed above, the latter two being in line with their putative regulatory nature.

### 3.3. CemR cysteine residues provide protein stability and coordinate Zn^2+^

As cysteine residues in proteins can play structural roles and may coordinate metal ions, we investigated the stability of each CemR protein variant and its ability to bind Zn²⁺.

We determined structural stability by analyzing protein thermal unfolding using nano-Differential Scanning Fluorimetry (nanoDSF) [53,54]. We measured the inflection point (IP), indicating the temperature at which 50% of the protein molecules unfolded. The IP of C_less_ and CD/DBD-CA muteins was approximately 7 °C lower than that of the wild-type protein and muteins in which RD-domain cysteine residues were replaced with alanine (Fig. 3A). Reducing conditions did not affect the stability of any CemR variant. Thus, the cysteine residues in the CD/DBD domain, which, according to AlphaFold3 structure prediction, are located in loops and/or unstructured regions (Fig. 1), may play a structure-stabilizing role. However, even when cysteine residues are preserved, the recombinant CD/DBD protein is unstable when isolated without the RD domain. Moreover, the C-terminal domain shows lower affinity and specificity for DNA than either the wild-type protein or a dimerizing GST-tagged variant of the CD/DBD domain (GST-CemR_CD/DBD_, Fig. S10). These data indicate that the CD/DBD domain requires the RD domain to maintain stability and structure and, consequently, DNA-binding activity.

**Fig. 3.**
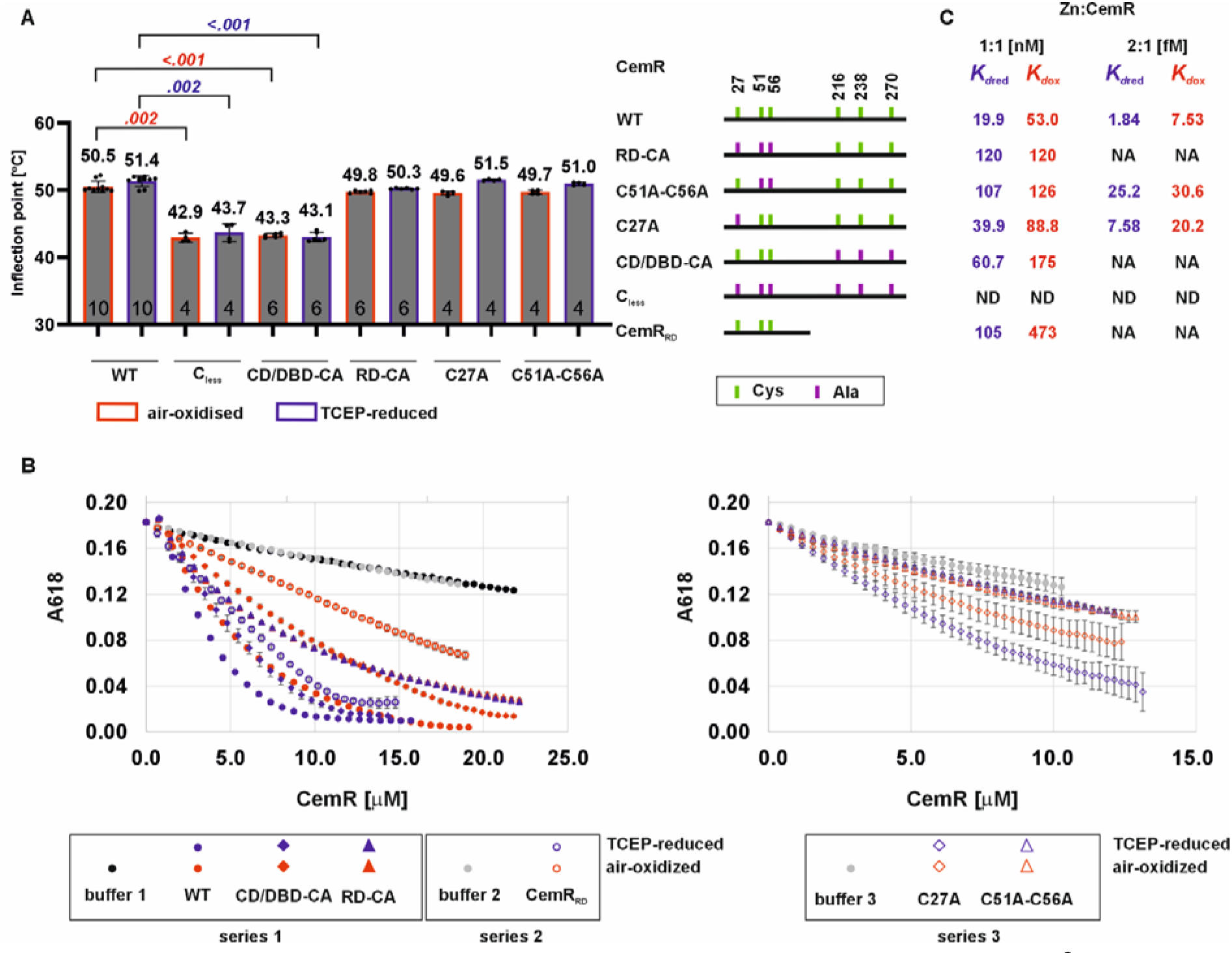
The role of cysteine residues in the stability of the CemR protein structure and Zn^2+^ binding. (**A**) Analysis of the thermal stability of the unreduced and TCEP-reduced wild-type CemR and muteins using nanoDSF. The temperature at which the stoichiometry of folded to unfolded protein molecules is 1:1 is determined by the inflection point value (IP). Analysis was carried out in at least two independent replicates, with two technical replicates in each. Data are shown as mean ±SD from n measurements; n for each protein variant is indicated at the bottom of the bar. Mann-Whitney test was used to determine the *P* value, with n measurements. *P*-values in blue and red indicate statistical significance between different muteins and WT protein under the same conditions (air-oxidized or TCEP-reduced); non-significant differences (P > .05) are not marked. (**B**) Analysis of Zn^2+^ binding to CemR wild-type and muteins. Reduced and non-reduced proteins were gradually added to a Zn^2+^-Zincon complex. The binding of Zn^2+^ to the proteins was analyzed by characteristic absorption decreases of the Zn^2+^-Zincon complex at 618 nm. The buffer without protein was used as a control in each series of analyses (series 1: WT, RD-CA, CD/DBD-CA; series 2: CemR_RD_; series 3: C27A, C51-C56). Data are shown as mean ± SD, with n = 3 independent replicates for all but the C51-C56 variant, for which only two replicates were used. (**C**) Dissociation constants determined from Zincon analysis (B). Assuming Zn^2+^: protein stoichiometry of 1:1 (Zn:CemR) *K*_dred_ and *K*_dox_ (both with M unit) denote dissociation constants of the reduced and air-oxidized proteins, respectively. Assuming Zn^2+^: protein stoichiometry of 2:1 (Zn_2_:CemR) *K*_d12red_ and *K*_d12ox_ (both with M^2^ unit) denote cumulative dissociation constants of the reduced and air-oxidized proteins, respectively (see Table S3 for numerical values of all dissociation constants). NA, not applicable because the CD/DBD domain or one domain’s cysteine residues are missing; ND, not determined.

The role of cysteine residues in Zn^2+^ binding was analyzed using the Zincon assay[39,55]. In this analysis, we monitored Zn^2+^-protein interaction by changes in absorbance at 618 nm of the Zn-Zincon complex resulting from the protein’s competitive binding of Zn^2+^. We previously demonstrated that the wild-type protein binds Zn^2+^ with increased affinity under reducing conditions, whereas the C_less_ variant of CemR does not bind zinc ions [39]. Therefore, cysteine residues are critical for Zn^2+^ binding. To more precisely analyze which cysteine residues in which domain are involved in Zn^2+^ binding, we used a series of air-oxidized and reduced CemR muteins. TCEP was used as a reducing agent because, in contrast to DTT or β-mercaptoethanol, it has a low affinity for Zn^2+^ [61]. Interestingly, our results indicate that cysteine residues in each domain engage in Zn^2+^ complexation by CemR (Fig. 3B). CemR_RD_ bound Zn^2+^ as an isolated domain, and both analyzed muteins (CemR_RD_ or CD/DBD-CA) bound Zn^2+^ in a redox-dependent manner. We observed no redox dependency in Zn^2+^ -RD-CA interactions. Zn^2+^ binding by the C27A mutein was redox-dependent, whereas the C51-C56 mutein interacted with Zn^2+^ similarly regardless of redox conditions (Fig. 3B). Thus, among the RD domain cysteine residues, C51 and C56, but not C27, were critical for redox-dependent activity. Since we were unable to obtain reliable data for the CemR_CD/DBD_ variant due to its low concentration and instability, it is impossible to determine whether CD/DBD cysteine residues participate in the formation of a second Zn^2+^ binding site independent of the RD domain or whether they cooperate, possibly allosterically, with cysteine residues of the RD to increase the affinity of the RD for Zn^2+^. No typical Zn^2+^-binding site can be predicted in CD/DBD, and AlphaFold low-probability prediction indicates H249 in the linker between the CD and DBD domains as putatively involved in Zn^2+^ coordination (Fig. S1). Thus, at this stage, it is impossible to unequivocally determine the affinity constant(s) for CemR interaction with Zn^2+^; however, for CemR_RD_, 1:1 stoichiometry is most likely. The dissociation constant of CemR_RD_ is estimated to be 4.73×10^-7^ M for the oxidized protein (*K*_dox_), decreasing to 1.05×10^-7^ M upon protein reduction (*K*_dred_) (Fig. 3C and Supplementary Table S3). Assuming 1:1 stoichiometry, WT protein affinity for Zn^2+^ decreases upon reduction from *K*_dox_ 5.30×10^-8^ M to *K*_dred_ 1.99×10^-8^ M. However, CemR may have two binding sites, with cumulative *K*_d12_ values of 7.53×10^-15^ M^2^ and 1.84×10^-15^ M^2^ for the oxidized (*K*_d12ox_) and reduced (*K*_d12red_) protein forms, respectively. However, each Zn^2+^ binding site can exhibit different affinities (*K*_d1_ ≠ *K*_d2_; note that only the cumulative *K*_d12_ value can be calculated from Zincon assay), which, together with redox-reactive interactions at the RD domain site, complicate the interpretation of the interactions and require further dedicated studies (see Discussion).

In conclusion, CemR contains at least one Zn^2+^ binding site in the RD domain, which binds Zn^2+^ even without the CD/DBD domain. Only this domain binds Zn^2+^ in a redox-dependent manner, with C51 and C56 being critical for redox-response-related Zn^2+^ - CemR interactions. Cysteine residues in the CD/DBD domain stabilize the CemR structure and are important for Zn^2+^ binding, with no difference between redox conditions; they either form a second Zn^2+^ binding site or collaborate with the RD domain to bind Zn^2+^.

### 3.4. CD/DBD-domain cysteine residues control redox-dependent CemR-DNA interactions

We previously demonstrated, using EMSA and SPR, that CemR interacts with DNA in a redox-dependent manner: non-reduced (air-oxidized) CemR binds DNA with lower affinity than TCEP-reduced CemR [39]. Cysteine residues play a role in redox-dependent DNA binding, because the C_less_ mutein was insensitive to redox reactions. However, since C_less_ CemR binds DNA, cysteine residues are not critical for direct DNA contact.

To characterize the role of cysteine residues in CemR redox-dependent DNA binding, we analyzed CemR variant activity under reducing (TCEP- or DTT-reduced), non-reducing (air-oxidized), or oxidizing (diamide-oxidized) conditions. We also investigated the reversibility of redox reactions depending on the presence or absence of cysteine residues. For this purpose, we first treated non-reduced CemR with diamide, and then reduced it with TCEP at a 5-fold excess over diamide (Materials and Methods). The CemR interaction with DNA was subsequently analyzed using SPR and/or EMSA.

The reduced WT CemR protein bound DNA with higher affinity and efficiency than the non-reduced protein (Fig. 4A), as we reported [39]. Diamide-oxidized WT CemR did not bind DNA (Fig. 4G); however, TCEP reduction of diamide-oxidized CemR restored its DNA-binding activity (Fig. 4G). The C_less_ protein bound to DNA under all redox conditions with similar affinity and efficiency, confirming that the redox reactions are specific to cysteine residues (Figs 4BH). Analysis of oxidized CD/DBD-CA interactions with DNA showed they were insensitive to DTT/TCEP reduction or diamide oxidation (Figs 4DJ), similar to the C_less_ protein variant. This suggested that cysteine residues in the CD/DBD-domain’s cysteine residues, but not in the RD domain, were responsible for redox-dependent regulation of CemR wild-type binding to DNA. Indeed, RD-CA mutein, in which cysteine residues at the CD/DBD domain were preserved, was sensitive to redox conditions: DTT reduction increased its DNA binding (Fig. 4C) while diamide oxidation inhibited its DNA binding activity (Fig. 4I). However, unlike in the WT protein, the oxidation was not fully reversible, because the subsequently reduced protein interacted with DNA with lower affinity than the TCEP-reduced CemR (Fig. 4I). These data suggested that either the RD domain’s cysteine residues were important for reducing oxidized CD/DBD domain cysteine residues, or they protected CD/DBD domain’s cysteine residues from excessive, irreversible oxidation. These results were confirmed by analysis of C27A and C51A-C56A muteins’ DNA binding. In SPR, these muteins bound DNA more strongly under reducing than oxidizing conditions (Fig. 4EF), similar to WT and RD-CA variants. In EMSA, diamide oxidation was nearly irreversible and inhibited mutein binding to CemR-DNA (Fig. 4KL), similar to what was observed with the RD-CA variant.

**Fig. 4.**
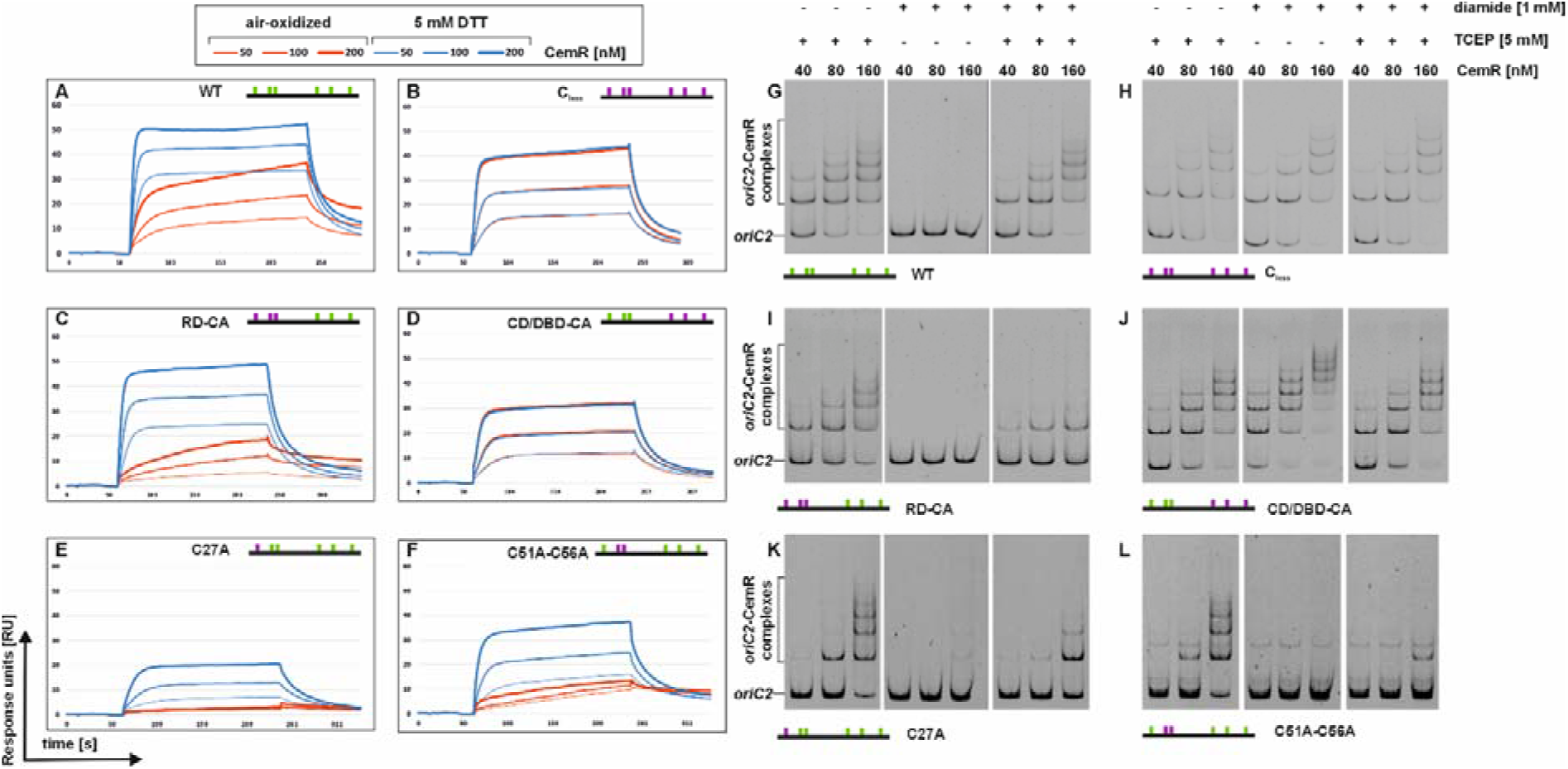
Influence of the redox state of CemR on its DNA-binding activity. (**A-F**): SPR analysis of interactions between CemR variants (50, 100, and 200 nM) and *oriC2* in the presence or absence of DTT. Protein concentrations are indicated by lines of varying thickness and color: red (CemR air-oxidized) or blue (CemR DTT-reduced). RU, response units. (**G-L)**: EMSA was performed using the FAM-labeled *oriC2* fragment, which was incubated with the indicated amounts of CemR wild-type and muteins in the presence or absence of TCEP or diamide. The results shown in panels G–L were obtained by separating samples of a given protein variant on a single polyacrylamide gel. Digital processing was applied equally across the entire image before it was divided into three subpanels representing the given conditions.

In conclusion, the CD/DBD domain’s cysteine residues directly mediate redox regulation of DNA binding by CemR. In contrast, the RD domain’s cysteine residues are important for the reversibility of this redox-dependent regulation. This analysis also confirms the interplay between the RD and CD/DBD domains, which may enable signal transduction between them.

### 3.5. RD-domain cysteine residues control Zn^2+^-redox-dependent CemR-DNA interactions

CemR interacted with Zn^2+^, and cysteine residues in each domain bound Zn^2+^ ([39] and Fig. 3BC). Moreover, in the presence of Zn^2+^, the DNA-binding activity of wild-type CemR, but not C_less_ CemR, was inhibited [39]. The inhibitory effect of Zn^2+^ on CemR-DNA interactions was particularly strong in the case of reduced protein. To further characterize the role of Zn^2+^ regulating CemR interaction with DNA, we analyzed how different CemR variants interact with DNA in the presence or absence of Zn^2+^ ions.

We confirmed that Zn^2+^ destabilizes the interaction of wild-type CemR with DNA, and that this effect is further increased in TCEP-reduced CemR (Fig. 5A). The presence of EDTA, regardless of whether it was added before or after incubation of CemR-*oriC2* with Zn^2+^, counteracted the inhibitory effect of zinc ions (Fig. S11A). This confirmed that destabilization is specific to Zn^2+^ as a divalent metal ion and that Zn^2+^-triggered destabilization of the CemR-*oriC2* complex is reversible (Figs 5A and S11A). Marginal Zn^2+^-dependent destabilization was observed for C_less_ CemR (Figs 5B and S11B), confirming that cysteine residues are involved in Zn^2+^-dependent regulation. Zn^2+^-destabilized RD-CA variant interactions with DNA, but TCEP reduction of RD-CA increased RD-CA binding to DNA, confirming the role of DC/DBD’s reduced cysteine residues in CemR DNA-binding activity. It also indicated that Zn^2+^-dependent inhibition of CemR binding to DNA may occur primarily in its RD domain. Indeed, Zn^2+^ strongly destabilized CD/DBD-DNA interactions, whereas Zn^2+^ did not affect binding by the non-reduced protein. Thus, coordination of Zn^2+^ by RD-domain cysteine residues in the reduced state inhibits protein interactions with DNA, although interaction with DNA is mediated by the CD/DBD domain. Residues C51-C56 are particularly involved in this regulation because replacing these residues with alanine, in contrast to C27, produces an effect similar to replacing all RD cysteine residues with alanine. Zn²⁺ inhibits the interaction of the C51A-C56A protein at a similar level in both redox states, and reduction of the protein slightly increases its binding to DNA, probably due to the reduction of the cysteine residues in the CD/DBD domain. The unreduced form of the C27A protein responds only slightly to the presence of Zn²⁺, while reduction of the protein, probably of the C51-C56 disulfide bond, increases the inhibitory effect of Zn²⁺, putatively by enabling Zn²⁺ coordination by the RD domain.

**Fig. 5.**
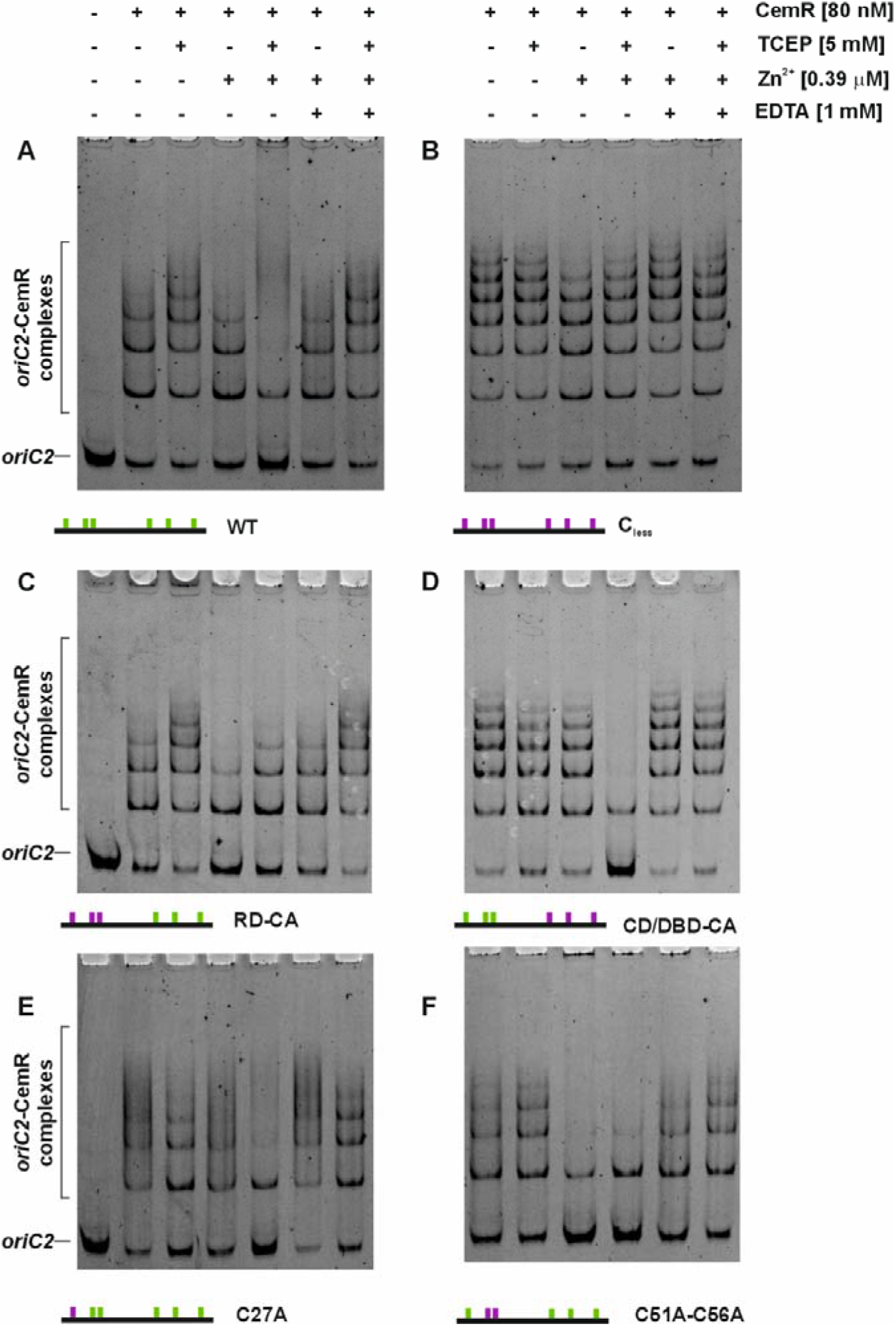
Influence of Zn^2+^ on CemR DNA-binding activity. EMSA was performed using the FAM-labeled *oriC2* fragment, incubated with the indicated amounts of CemR protein (wild-type and muteins) in the presence or absence of Zn^2+^. Digital processing was applied equally across all images, including controls.

In summary, the RD domain regulates CemR binding activity in response to redox state and Zn²⁺ presence. Zn^2+^ coordination, likely involving the reduced residues C51 and C56, inhibits CemR interaction with DNA.

### 3.6. The C51-C56 residues are important for regulating gene expression in vivo

Our previous studies on CemR indicated that *gluP* and *fecA3* (HP1174 and HP1400 in *H. pylori* 26695, respectively) are directly controlled by CemR [39,42]. These genes are also controlled by other pleiotropic regulators. *gluP,* which encodes an efflux pump of the major facilitator superfamily (MFS) involved in glucose transport and antibiotic resistance [62,63], is possibly controlled by Fur[64] and, in biofilms, by SpoT via the σ54-dependent pathway [62]. *GluP* expression initially increases at low pH but decreases after prolonged acid stress [65,66]. *fecA3,* whose transcription is phase-growth-dependent, peaking in the logarithmic phase and declining in the stationary phase [67,68], is primarily controlled by NikR [68,69] and, in the stationary phase, possibly additionally by Fur [67,68]. In our previous studies, CemR repressed *gluP* and *fecA3* under microaerobic conditions, and these genes were derepressed under aerobic stress. This control was lost in the deletion ΔCemR *H. pylori* mutant, and *gluP* and *fecA3* were upregulated compared to the unstressed WT N6 strain, regardless of redox conditions [39]. In the cysteine-less ΔCys strain, *gluP* and *fecA3* were expressed at higher levels than in the WT strain under both redox conditions, indicating that cysteine residues regulate CemR transcriptional activity. Our *in vitro* research presented in this work indicated that the RD domain cysteine residues play an important role in regulating CemR activity. Therefore, we prepared a mutant strain in which we replaced a single C27 residue and a pair of C51 and C56 residues with alanine (C27A and C51A-C56A, respectively; see Materials and Methods). We then examined *gluP* and *fecA3* expression under microaerobic and aerobic conditions.

*gluP,* as in previous studies, was derepressed in the WT strain under aerobic conditions compared with microaerobic growth (Fold change FC of 3.0±0.9), and its transcription increased two-to three-fold in ΔCemR regardless of redox conditions (Fig. 6A). In the ΔCys strain, *gluP* was expressed at a similar level as in the WT strain under microaerobic conditions (Fig. 6), unlike previously observed [39], possibly due to differences in experimental conditions between analyses and multilevel regulation. Nonetheless, *gluP* transcription in the ΔCys strain was independent of redox conditions in two independent sets of experiments (Fig. 6 and [39]), indicating that cysteine residues of CemR are important for redox-dependent *gluP* transcriptional response. CemR C27A protein controlled *gluP* expression similarly to the WT protein, suggesting that C27 is not critical for CemR control over *gluP*. In the C51A-C56A CemR mutant, *gluP* transcription was repressed approximately twofold under both redox conditions, indicating that these two cysteine residues are important for *gluP* transcription levels and its redox-dependent control.

**Fig. 6.**
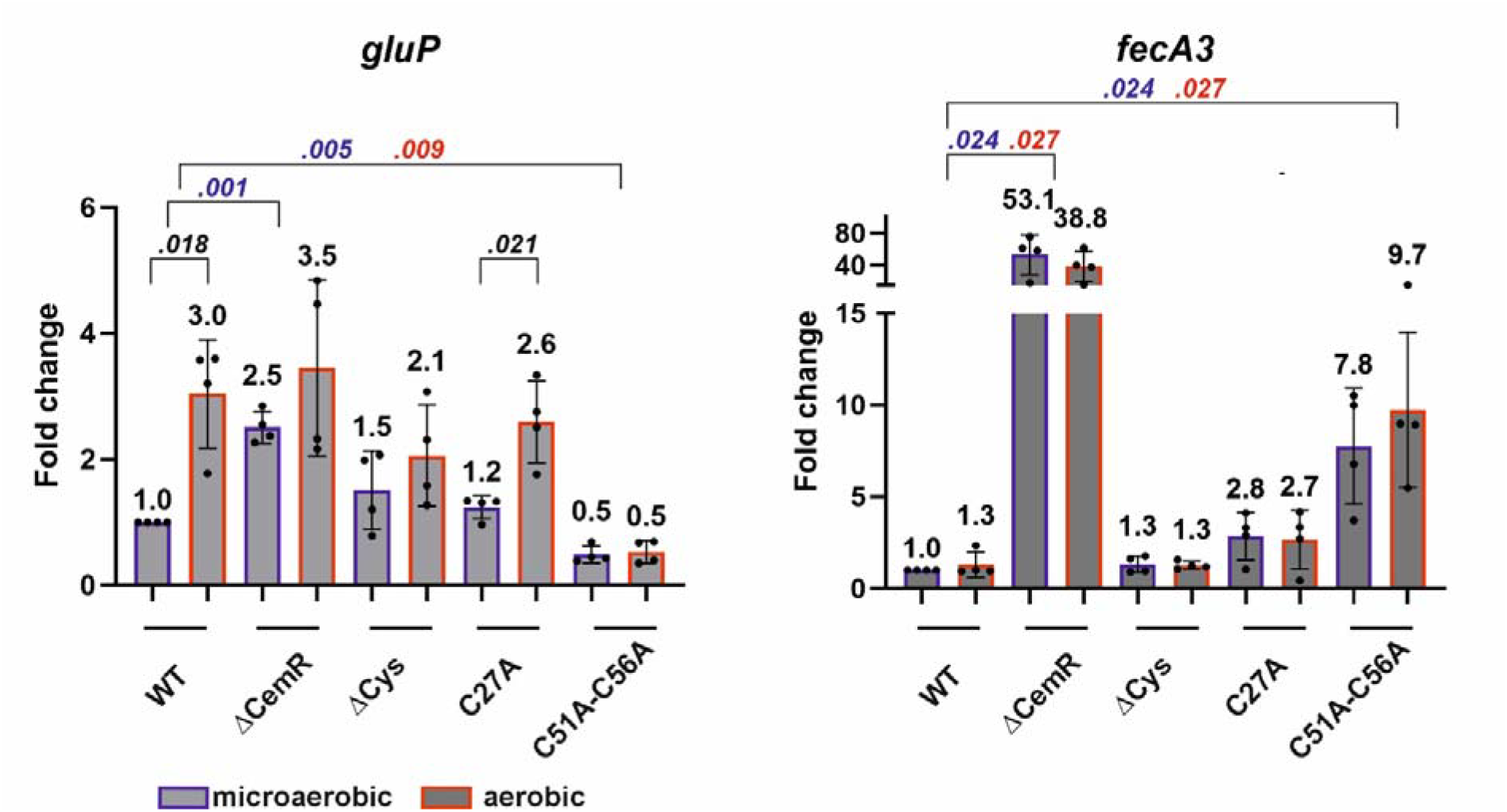
RT-qPCR analysis of *gluP* and *fecA3* transcription in *H. pylori* N6 cultured under microaerobic and aerobic conditions. Results are presented as fold changes relative to the WT strain under microaerobic conditions. Data are shown as mean ± SD from 4 biological replicates. A two-tailed Student’s t-test (Welch’s test) determined the *P*-value. P-values in blue and red present statistical significance between transcription in different strains under microaerobic and aerobic conditions, respectively; *P*-values in black present statistical significance under transcription in the same strain between microaerobic and aerobic conditions; only significant P-values (P> .05) are shown.

For *fecA3* transcription, we did not observe redox-dependent transcriptional regulation in any strain (Fig. 6). The result for the WT (FC 1.3±0.7) differs from previous studies [39]; however, the difference previously reported was also relatively small (FC 2.0±0.4) [39]. Nonetheless, this study confirmed a marked increase in *fecA3* transcription in the ΔCemR mutant. The *fecA3* gene was slightly induced in the C27 mutant regardless of redox conditions; however, the difference was not statistically significant (P> .05). In a CemR C51A-C56A mutant, *fecA3* transcription was further induced, approximately 8-fold compared to the WT strain, indicating a stronger influence of the C51-C56 residues *fecA3* transcription than that of C27.

In summary, despite differences in the regulation of the *gluP* and *fecA3* genes across two independent sets of experiments likely arising from complex regulatory mechanisms mediated by CemR, Fur, and NikR—we have unequivocally demonstrated that the CemR C51–C56 residues play a significant role in regulating transcription.

## 4. Discussion

The systems regulating H. pylori’s response to stresses are very limited [18,70]. CemR is the least typical of the 13 H. pylori proteins identified to regulate gene transcription in response to intra- or extracellular signals [18] (see below) and one of the few characterized bacterial redox switches that evolved from two-component systems such as Streptococcus pneumoniae RitR [3,71], Staphylococcus aureus AgrA [72], or Vibrio cholerae ArcA [73]. Although CemR consists of typical modules found in response regulators (RR) of the TCS systems, namely the N-terminal signal-receiving domain (RD), the central domain (CD), and the C-terminal DNA-binding domain (DBD), only the DBD with its HTH motif exhibits the typical features. The RD domain is not only classified as an ALR domain but is also redox- and Zn -regulated, which is atypical for *RR. The CD domain, also controlled by redox and Zn²⁺, is not homologous to any domains described so far. Furthermore, CemR activity appears to be largely controlled by allosteric interactions between the REC, CD, and DBD domains, possibly mediated by a signaling* α *helix [74] and unstructured regions, which together indicate that this protein is characteristic only of the class of* **Campylobacteria**.

### 4.1. CemR domains shape protein regulation

The RD domain tertiary structure is typical for RR proteins (Figs S2 and S3). However, its sequence is altered at key sites important for phosphorylation-mediated signal transduction, including a phosphate-binding aspartic acid residue replaced by serine at position 47 (Fig. 1, Figs S2 and S3). In contrast, RD contains three cysteine residues, of which C51-C56 form an intramolecular disulfide bond in a recombinant, non-reduced protein purified under aerobic conditions (CemR_S-S_). The redox state of C51-C56 does not affect CemR’s DNA-binding activity per se; however, it affects CemR’s affinity for Zn^2+^, which in turn controls CemR’s affinity for DNA. In particular, CemR_S-S_ binds Zn^2+^, thereby reducing its affinity for DNA. Reduction of the S−S bond increases CemR’s affinity for Zn^2+^ significantly, further reducing CemR’s DNA affinity. Thus, conditions that affect the redox state of cysteine residues together with Zn^2+^ levels are critical factors regulating CemR-DNA interactions. No protein motif typically associated with Zn^2+^ binding is identified in *H. pylori* CemR, neither in RD nor in the CD/DBD domain (see Discussion below). Nonetheless, since the C_less_ protein does not bind Zn^2+^ [39], C51 and C56 are critical either for direct Zn^2+^ - CemR interactions or for shaping the CemR structure required for Zn^2+^ metalation (Fig. 5). Although the exact mechanism of how C51 and C56 redox state affects RD domain structure is unknown, it should be noted that these residues are located at the flexible region between η1 and α2, close to the region at which a phosphoacceptor aspartic acid residue is located in typical REC domains (Figs. S2 and S3). It suggests that S−S bond formation/reduction and Zn^2+^ binding may play roles similar to phosphorylation, triggering structural changes in the REC domain that are eventually, and possibly allosterically, passed to the CD/DBD domain. However, structural changes may be less typical than in bacterial REC domains, because a few conserved amino acids involved in these structural changes are replaced in CemR by amino acids with different biochemical properties (Figs S2 and S3).

An intriguing question that remains unanswered is the role of C27 in RD domain regulation. Our studies indicate that, unlike C51-C56 of RD and possibly C216 and C270 of the CD/DBD domains, C27 is not oxidized under non-reducing conditions in vitro. Its role in CemR regulation in vivo is less prominent than that of C51-C56 (Fig. 6). Yet, CemR C27 is the sole conserved cysteine residue in all CemR-family proteins across Campylobacterial species, suggesting its functional or structural importance under conditions yet to be uncovered or for promoters with complex structure such as those controlled by multiple regulators. C51-C56 are conserved only in Helicobacter species but are absent in closely related species such as Campylobacter jejuni or Arcobacter butzleri (Fig. S2), raising the question of how these regulators are controlled.

Although RD and DBD domains are homologous to those of the NtrC family regulators AtoC and RocR, respectively, the CD domain, unlike that of typical NtrC regulators, lacks motifs critical for AAA+ domain activity and the conserved GAFTGA sequence motif. These features are required for NtrC-like regulators’ oligomerization and interaction with the σ54 factor, respectively, thereby resulting in transcriptional control of gene expression. Moreover, the CD domain has no homologs in protein databases outside the class *Campylobacteria*. Both domains, CD and DBD, contain large unstructured regions in which C216, C238, and C270 residues are located between α11-β9, β10-η4 , and α13-α14, respectively; the latter cysteine residue lies at the border of the HTH motif. These cysteine residues, including the redox-sensitive C216 and C270, are not directly involved in DNA binding but instead play a structural role, increasing CemR affinity for DNA when reduced. They also engage in redox-independent Zn^2+^ binding, which partially inhibits CemR’s affinity for DNA, especially under non-reducing conditions. This apparent contradiction between the redox-independent interaction with Zn^2+^ and the redox-dependent interaction with DNA likely points to allosteric signal transduction from the CD domain to the DBD, altering the spatial arrangement of the HTH motif and affecting DNA binding. Thus, CD/DBD domains are the second protein region to receive the redox signal, thereby triggering the protein’s transcriptional response.

The RD and CD/DBD domains are connected by a long helix, which likely mediates signaling between the two [75]. Such helices are typical in the NtrC family of regulatory proteins (e.g., *A. aeolicus* NtrC1 [76], *Brucella abortus* NtrX [77] or *Sinorhizobium meliloti* DtcA [78]). The length of the helix in CemR correlates with the so-called “negative type” of NtrC regulation, in which the RD in an off state (e.g., unphosphorylated) interacts with the CD domain and blocks AAA+ oligomerization and interaction with factor σ^54^ of the RNA polymerase holoenzyme (RNAPH). Activation of the RD domain releases the AAA+ domain, allowing oligomerization, protein activation, and interactions with σ^54^-RNAPH [75,79]. Signaling helices also occur in less-typical redox-controlled regulators, such as the non-TCS *Myxococcus xanthus* PexR[80] and the *Streptococcus pneumoniae* TCS-derived non-NtrC-like RitR[3]. In PexR, the signaling helix is part of the N-terminal signal-perceiving GAF domain linked to the NtrC-like CD domain; a long unstructured region connects the C-terminal DBD domain to the CD domain. GAF binds either Zn^2+^ or nonheme Fe^2+^ *via* C122, H102, H120, and D140. Upon oxidation, the metal ion is lost, activating PexR and inducing gene transcription. However, neither the exact role of C122 in GAF activity control, nor any of the other six PexR cysteine residues have been characterized yet [80]. RitR is an orphan atypical response regulator composed of an ALR-type REC domain and an OmpR/PhoB DNA-binding domain connected by a long α helix [3]. At the C-terminus, this signaling helix contains a redox-regulated C128 residue that oxidized under elevated oxygen levels, forming a disulfide bond between RitR protomers in the dimer. This 2-Cys interdomain dimerization leads to helix unraveling and DBD domain swapping, thereby unblocking the DBD domains and allowing them to bind DNA. It was proposed that interdomain α-helix disulfide-bond signaling may be a more common mechanism, also found in *Campylobacterota* [81]. However, this is not the case for *H. pylori* CemR and its homologs, in which cysteine residues are located outside the domain-connecting α helix.

The mechanism of signal transduction between CemR domains and other transcriptional subunits is unknown, especially since information on CemR CD domain activity is scarce. Studies of that domain were hindered by the instability of the purified recombinant CemR_CD/DBD_ protein and by its loss of DNA-binding activity, which can be partially regained when the protein is fused to a dimerizing GST protein. Thus, either interaction with the RD domain or dimerization by RD is required to maintain the CD/DBD protein structure and function. GST may partially mimic both features, explaining why GST-CD/DBD binds DNA more strongly than CemR_CD/DBD_. Moreover, the RD domain must contain intact cysteine residues for efficient, reversible, N-terminal cysteine-dependent redox regulation of CemR DNA binding. When diamide oxidizes CD/DBD cysteine residues, thereby precluding DNA interaction, they can be efficiently reduced and regain DNA-binding activity only when the RD domain contains cysteine residues; i.e., the domains must interact, either directly or allosterically.

### 4.2. CemR and zinc homeostasis

Zinc homeostasis in *H. pylori*, unlike nickel and iron, remains poorly characterized [82,83]. Zn^2+^ is an essential trace element and a chemoattractant for *H. pylori*, and it is critical for bacterial physiology and virulence [82–88]. *H. pylori* copes with an environment that is occasionally poor or rich in Zn^2+^, sequestered by the host or supplied by the host diet, respectively [83]. No zinc uptake system has been identified, whereas three efflux systems have been described [82,83]. Yet, no Zn^2+^-controlled transcription regulator has been identified in *H. pylori* beyond Fur, a Zur-family regulator that is primarily controlled by iron levels but responds to Zn^2+^ levels to some extent [89,90]. The overall cellular concentration of total Zn^2+^ in bacterial cells is estimated in the submillimolar range, but the “free” (frequently called labile) Zn^2+^ concentration is estimated in the pico-to femtomolar ranges, because uncomplexed Zn^2+^ is toxic to biomolecules [91–93]. Consequently, femtomolar Zn^2+^ concentrations trigger transcription of Zn^2+^ uptake or efflux systems . Thus, Zn^2+^ is either bound to proteins (e.g., enzymes or metal-storage proteins) or to low-molecular-weight molecules such as GSH or bacillithiol (BSH), often synthesized by gram-negative or gram-positive bacteria, respectively [91,93,94]. The concentration of “free” Zn^2+^ in *H. pylori* cells is unknown. *H. pylori* lacks a glutaredoxin-mediated redox regulation system utilizing GSH, as well as GSH itself [95,96], which challenges not only redox but also zinc homeostasis. Moreover, Zn^2+^-storage or chaperone proteins have not been sufficiently investigated. Histidine-rich protein Hpn interacts with Zn^2+^, but Hpn’s affinity for Zn^2+^ is lower than for Ni^2+^ and Cu^2+^ [97]. Thus, the occasional fluctuations in the “free” Zn^2+^ level may be more frequent in *H. pylori*. It is tempting to speculate that CemR also helps maintain zinc homeostasis in *H. pylori*.

Our studies did not allow us to unequivocally determine the exact number of Zn^2+^-binding sites and, consequently, the exact affinity of CemR for Zn^2+^ ions at specific sites, which is essential for establishing whether Zn^2+^-dependent regulation is significant at physiologically sufficient Zn^2+^ levels or at elevated/decreased concentrations. No typical Zn^2+^ binding site can be identified in any CemR domain. AlphaFold3 predicted that surface-exposed H18 of the RD domain and H249 of the CD/DBD domain linker are involved in Zn^2+^ binding. Surface-exposed histidine residues have been identified as critical for Zn^2+^ binding in some proteins, such as H36 in the Zur protein [98]; H249 is predicted to be partially solvent-accessible and to create a binding site together with H249 of the second protomer of the dimer. However, these predictions were not experimentally validated. Redox regulation of each domain complicates studies and interpretation of the results by influencing CemR’s affinity for Zn^2+^, and the instability of the isolated CD/CBD domain further complicates them. If CemR binds only one Zn^2+^ ion, the nanomolar *K*_d_ values suggest it responds to relatively high “free” Zn^2+^ concentrations, possibly when extracellular Zn^2+^ levels exceed homeostatic levels. If we assume that CemR binds two Zn^2+^ ions, the average nanomolar-range affinity may not reflect the actual metalation status. It is more likely that one Zn^2+^ binding site is tighter (for example, picomolar or even lower) than the second one (being in the nanomolar range), allowing sequential Zn^2+^ binding depending on cellular “free” Zn^2+^ status. A consequence of the two regions’ differing affinities for Zn^2+^ is a broader response across a wider range of Zn^2+^ concentrations, possibly below and/or above homeostatic levels, depending on the gradual or collaborative saturation of CemR’s Zn^2+^ binding sites. Such a scenario was recently described for multiple Cys-rich Zn^2+^-binding sites in metallothioneins from different domains of life, proving that one binding site interacts with other ones by allostery and a hydrogen-bond network [99,100]. For example, it has recently been discovered that the *Streptomyces coelicolor* Zur protein has, in addition to previously known one structural site and two regulatory sites that bind Zn^2+^, the third regulatory site that also binds Zn^2+^. These sites likely differ by less than 20-fold in affinity within the femtomolar range per site, allowing modulation of Zn^2+^ occupancy at Zur binding sites. Consequently, it modulates Zur’s affinity for DNA in response to Zn^2+^ saturation, thereby enabling discrimination between promoters activated at high or low Zn^2+^ concentrations and allowing Zur to act as an inducer or a repressor [98]. Some transcriptional regulators, such as *Staphylococcus aureus* CzrA, exhibit negative allosteric regulation of Zn^2+^ affinity to distinct Zn^2+^ binding sites, also discriminating promoters based on Zn^2+^ concentration [101]. CemR acts as a repressor or activator of genes under both microaerobic conditions and oxidative stress, but the mechanism enabling CemR to discriminate promoters and trigger positive or negative transcriptional responses remains unknown. It is tempting to speculate that CemR binds Zn^2+^ at two binding sites with different affinities or allosterically modulates the affinity of a single Zn^2+^ binding site, which, together with redox regulation, provides a comprehensive response to the cell’s redox conditions and possibly also to “free” Zn^2+^ concentration (Fig. 7). It should be noted that upon oxidative stress, Zn^2+^ is released from the proteins due to cysteine oxidation, and consequently the concentration of “free” Zn^2+^ increases in the cells. Interestingly, some genes regulated by CemR under oxidative stress are similarly affected by near-toxic Zn^2+^ levels [102]. These include the directly CemR-controlled *gluP, comB8*, and *comB9* (HP0038 and HP0039, components of the ComB DNA uptake system), and *hemH* ferrochelatase (HP0376), which are derepressed under aerobic stress and increased Zn^2+^ levels, while HP1192, encoding a secreted protein involved in *H. pylori* motility, is repressed under stress [39,42,102]. Approximately 30% of genes indirectly controlled by CemR under oxidative stress are similarly induced or repressed by toxic levels of Zn^2+^. In contrast, only five of 136 genes (excluding RNA genes) are regulated in the opposite direction. In our studies, we mainly monitored DNA-affinity changes of CemR; *in vivo,* however, CemR binds to its targets regardless of redox conditions [42]. Thus, regulatory effects may involve structural changes in the CemR nucleoprotein complex that affect either DNA promoter structure, and thereby, other DNA-protein binding, or CemR interactions with other transcription machinery subunits such as RNA polymerase. Further investigation is needed to determine whether changes in Zn^2+^ levels are detected directly by CemR, or whether excess Zn^2+^ induces stress in the cell – including oxidative stress – and whether this oxidative stress specifically triggers CemR’s response.

**Figure 7.**
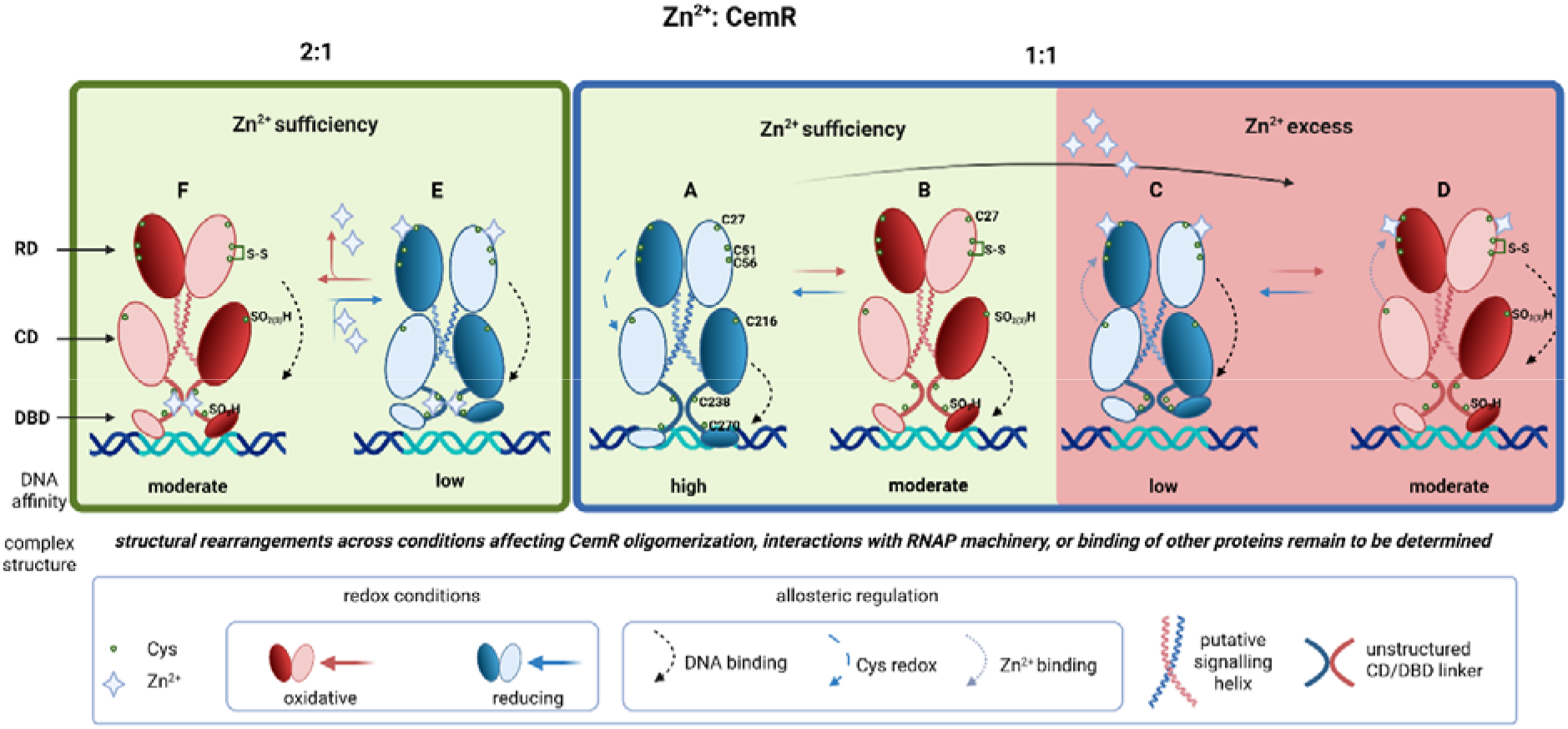
A model of CemR activity regulation. In a 1:1 Zn^2+^:CemR model, the protein does not interact with Zn^2+^ at physiological, putatively femtomolar Zn^2+^ levels because of its low affinity. (A) Under reducing conditions, apo-CemR has a high affinity for DNA, forming a specific apo-CemR-DNA complex. (B) Under oxidizing conditions, cysteine residues are oxidized, reducing apo-CemR affinity for DNA and reorganizing the nucleoprotein complex, specifically due to oxidation of the CD and CD-DBD linker. Oxidized cysteine residues in the CD/DBD can be reduced, possibly involving cysteine residues in the RD domain. (C) When Zn^2+^ levels rise under reducing conditions, CemR binds metal, particularly via the REC domain, and holo-CemR significantly alters its affinity for DNA or restructures the nucleoprotein complex. (D) Under oxidizing conditions, holo-CemR’s affinity for DNA or a rearrangement of the complex resembles that of oxidized apo-CemR. In the 2:1 Zn^2+^: CemR model. (E) CemR shows a high affinity for Zn²⁺ under optimal physiological conditions (reducing cytoplasmic conditions, sufficient Zn^2+^), and both domains may bind Zn^2+^. Holo-CemR interacts with DNA with very low affinity, forming a complex with a specific structure. (F) Under oxidative stress, CemR undergoes oxidation, and residues C51–C56 form a disulfide bridge. The REC domain’s affinity for Zn^2+^ decreases, which allosterically increases apo-CemR’s affinity for DNA or restructures the nucleoprotein complex, changing the transcriptional response. More details are in the text. Created in https://BioRender.com

### 4.3. CemR integrates into H. pylori signal transduction system

CemR’s unusual structure and multimodal regulation may stem from its integration into the gene transcription regulatory circuit, mediated by a few, mostly pleiotropic regulators that provide a broad spectrum of regulation despite a limited repertoire of proteins [18,70,103]. In particular, an atypical structure may reduce stochastic crosstalk with components of other TCS systems or enable such crosstalk only under specific conditions [104]. *H. pylori* encodes three complete phosphorylation-controlled TCS systems, namely ArsRS (HP0166/HP1065), CrdRS (HP1365/HP1364), and FlgRS (HP0703/HP0244), which regulate the responses to acid stress, nitrosative stress, and flagellar biosynthesis, respectively. HsrA (HP1043), similarly to CemR (HP1021), is an orphan atypical response regulator. HsrA is an essential protein that, despite years of research, responds to still unknown stimuli ([18] and references therein). Structurally and mechanistically, HsrA, ArsR, and CrdR belong to the OmpR family of proteins [105], which usually control transcription by interacting with DNA at the gene promoter region and core subunits of the RNA polymerase holoenzyme (RNAPH) or the housekeeping σ^70^ factor (σ^80^ in *H. pylori*, HP0088) of RNAPH[106–108]. FlgR belongs to the NtrC family of regulators [105], also called bacterial enhancer-binding protein (bEBP), which typically binds DNA via the C-terminal domain and activates transcription by interacting with the σ^54^ factor (*H. pylori* HP0714) of RNAPH[79]. However, FlgR is an atypical regulator lacking a DNA-binding domain [109]. This means it lacks the domain that confers specificity in regulating a particular gene’s transcription, and to provide that specificity, other proteins had to be devoid of the NtrC-like domains. This likely explains why the CD domain of CemR, despite being flanked by RD and DBD domains typical of NtrC family proteins (AtoC and RocR, respectively), has been replaced by a different domain not associated with NtrC. Interestingly, *C. jejuni* FlgR contains a C-terminal domain lacking a DNA-binding motif, which prevents *in vivo* crosstalk and modification by acetyl phosphate, limiting phosphorylation specificity to FlgS-FlgR transfer [110,111]. *H. pylori* also uses two metalloregulators, Fur and NikR, primarily controlled by the levels of iron and nickel, respectively. However, redox conditions have also been suggested to regulate Fur [112], while NikR has been demonstrated to respond to acidity [113,114] and oxidative stress [115,116] regardless of the nickel presence. However, the response to acid stress, critical for *H. pylori* survival, is controlled not only by ArsRS [65], but also by FlgS (FlgR-independently)[117], and NikR [113,114]. Oxidative stress is controlled by CemR [39,42], but also Fur [118,119], HsrA [120], NikR [115], ArsR [65,121] and CsrA [122]. These overlapping activities of different regulators in response to a given stress, but responding to stimuli within different ranges (e.g., highly or mildly acidic pH or variable ROS factors, in the presence or absence of metal ions), possibly enable *H. pylori* to manage a broad range of stresses using a limited regulatory resource. However, this approach risks crosstalk between regulators, such as phosphorylation of response regulators by non-cognate KHs, which was previously considered disadvantageous [104,123,124], or nonspecific interactions between proteins with similar domains. Therefore, changing the regulatory mechanism of CemR from typical phosphorylation of the REC domain to a metal-Cys redox switch minimizes crosstalk with other TCS systems and eliminates the need for encoding an additional, CemR-dedicated kinase. Interestingly, TCS-derived response regulators controlled via a combined metal-thiol redox mechanism are atypical, and we found no examples of such response regulators, although this regulation occurs in HK (e.g., the α-proteobacterial RegB kinase [125]). It will be interesting to investigate further the molecular mechanism regulating CemR activity, including both metal-redox signal transduction and the possibility of regulation via the increasingly recognized cross-activation by kinases other than HK [21], or phosphorylation or acetylation by low-molecular-weight substrates derived from metabolism. Such mechanism has not yet been investigated in *H. pylori*. These post-translational modifications are especially interesting because CemR in *H. pylori* and other related *Campylobacter* species play a major role in regulating central carbon metabolism [38]. Dual redox and phosphorylation regulatory systems have been reported for response regulators such as RitR, *Staphylococcus aureus* AgrA, or *Vibrio cholerae* ArcA. RitR’s transcriptional activity depends on the StkP kinase-PhpP phosphatase system beyond previously mentioned redox control [71]. In ArcA, primarily controlled by phosphorylation by ArcB HK[126], redox control mediated by an intramolecular disulfide between C173 and C233 has recently been identified [73]. AgrA is regulated by a complex redox mechanism that leads to intramolecular disulfide bond formation (C199-288) and modification of the C6 and/or C199 cysteine residues coenzyme A [72,127], as well as by post-translational modifications via the classical TCS pathway [128]. Further studies on co-regulation in *H. pylori*, which has a limited repertoire of regulatory proteins, are needed to unravel the superior adaptability of this organism in its hostile niche.

## Data availability

The mass spectrometry data have been deposited in the PRIDE database (Project accession: PXD085775).

## Acknowledgments

We thank Mr. Fabian Pikuła for his contributions to this research: (1) adapting differential thiol-trapping protocols for CemR analysis, particularly IA/AMS labeling, and performing these analyses (Fig. 2); (2) conducting preliminary LC–MS/MS analyses under the supervision of K.K.; (3) performing zincon analyses of WT, RD-CA, CD/DBD-CA, and CemR_RD_ (Fig. 3); and (4) establishing conditions for reversible regulation of redox activity and testing selected proteins under those conditions (Fig. 4G–J).

The purchase of M-Class Acquity-Synapt XS LC-MS and Prometheus Panta systems was financially supported by the “Excellence Initiative – Research University” program for the University of Wroclaw (2020–2026). Biophysics and Bioanalytics Facility (RRID:SCR_021630) was funded as part of the IIMCB IN-MOL-CELL Infrastructure by the European Union – NextGenerationEU under the National Recovery and Resilience Plan. IN-MOL-CELL Infrastructure was also funded by the European Union under Horizon Europe (Project 101059801 -RACE) and by RACE-PRIME project carried out within the IRAP programme of the Foundation for Polish Science co-financed by the European Union under the European Funds for Smart Economy 2021-2027 (FENG). We thank NanoTemper Technologies for providing access to the Prometheus NT. 48. device for testing.

## Author contributions

Conceptualization: A.Z.P.; Methodology: A.Z.P., K.K., A.K.; Software: M.T.; Validation: A.Z.P.; Formal analysis: A.Z.P., M.T., A.K.; Investigation: J.M., P. Sz., M. N., K.K., K.S., M.T.; Resources: A.Z.P., K.S., A.Cz., M.T., A.K., K.K.; Data curation: A.Z.P., M.T.; Writing – Original Draft: A.Z.P., J.M., M.T.; Writing – Review and Editing: A.Z.P., J.M., M.T., A.Cz, A.K., K.S., P. Sz., M.N. K.K.; Visualization: A.Z.P., J.M., M.T., K.S., A.K.; Supervision A.Z.P., A.K.; Project administration and Funding acquisition: A.Z.P.

