## Supplementary Material, Tables and Figures for "Zinc- and redox-dependent allosteric regulation of the *Helicobacter pylori* CemR transcription factor"

**This PDF file includes:**

**Supplementary Material and Methods**

**Supplementary Tables 1 to 3**

- Tab. S1. Strains, plasmids, and proteins used in this study.
- Tab. S2. Primers used in this study
- Tab. S3. Dissociation constants of $Zn^{2+}$ complexes with CemR variants at various oxidation states.

**Supplementary References**

**Supplementary Figures 1 to 11**

**Supplementary Material and Methods**

**Construction of the *H. pylori* mutant strains**

To construct a vector for simultaneous exchange of the codons C51 (TGC) and C56 (TGC) for A51 (GCC) and A56 (GCC) in HP1021 on the *H. pylori* chromosome, the HP1021 gene was amplified using the P22/P27 and P21/P28 primer pairs and the pUC18HP1021com template. The fragments were cloned by Gibson Assembly[1] into SmaI-digested pUC19, generating pUC19/HP1021_C51A-C56A.

To construct a vector for exchange of the codon C27 to A27 (TGC→GCT) in HP1021 on the *H. pylori* chromosome, the pTZ57R/TΔHP1021 vector was amplified using the phosphorylated P25/P26 primer pair, and religated, introducing new BglII and XhoI restriction sites into pTZ57R/TΔHP1021 and generating pTZ57R/TΔHP1021_1. The HP1021 C27A gene was excised from pET28StrepCemRC27A using BglII and XhoI and cloned into pTZ57R/TΔHP1021_1, generating the pTZ57R/THP1021_C27A vector used for allelic exchange of HP1021 *via* homologous recombination.

*H. pylori* mutant strains were constructed using a homologous recombination approach as described by Ge and Taylor[2]. Briefly, *H. pylori* cells were plated from the stock to CBA-B plates and incubated under microaerobic conditions for 24 h. Next, cells were plated on CBA-B plates and incubated for approximately 5 h to form a micro-lawn 10 mm in diameter. Then, the micro-lawn was spotted with 10 µg of purified recombinant plasmids and grown under microaerobic conditions. After 24 h of growth, the culture was plated onto a selective medium and incubated for 5 days to obtain single colonies of transformants. The allelic exchange was verified by PCR using the P23-P24 primers, which are homologous to the chromosomal regions flanking the designed recombination sites, and by sequencing the PCR products with the P15-P34 primer pair.

**Construction of plasmids expressing recombinant wild-type and mutant HP1021 variants**

pET28Strep, which produces strep-tagged proteins at the N-terminus, was chosen as an expression vector for the synthesis of the wild-type CemR and its cysteine-to-alanine variants (Supplementary Table S1). The plasmids used for the expression of wild-type (pET28StrepHP1021), RD-CA (pET28StrepHP1021NΔCys), CD/CBD-CA (pET28StrepHP1021CΔCys), and $C_{less}$ (pET28StrepHP1021ΔCys) CemR variants were constructed before[3].

To construct vectors for the synthsisis of $CemR_{RD}$ and $CemR_{CD/DBD}$, the HP1021 was amplified with primer pairs P15/P16 and P17/P18, respectively, using pET28StrepHP1021 DNA as a template. The PCR products were gel-purified, digested with BamHI/XhoI, and cloned into the BamHI/XhoI sites of pET28Strep to generate $pET28StrepCemR_{RD}$ and $pET28StrepCemR_{CD/DBD}$, respectively. To construct a vector for the synthsisis of GST-tagged C-terminal domain of CemR, the HP1021 was excised from $pETStrepCemR_{CD/DBD}$ using BamHI/XhoI and cloned into the same sites of pGEX-6P-2 to generate $pGexCemR_{CD/DBD}$.

To construct a vector for the synthsisis of CemR C27A, in which the C27 codon was exchanged for the A27 (TGC→GCT), the HP1021 gene was amplified with primer pairs P15/P20 and P19/P18 using pETStrepHP1021 DNA as a template. The PCR products were gel-purified and used as templates in an overlapping extension PCR (OE-PCR[4] ) reaction with the P15/ P18 primer pair. The PCR product was purified, digested with BamHI/XhoI, and cloned into the BamHI/XhoI sites of pET28Strep to generate pETStrepCemRC27A (Supplementary Table S1). To construct a vector for the synthsisis of CemR C51A-C56A, in which the codons C51A and C56A were exchanged for A51 and A56 (each TGC→GCC), the HP1021 gene was amplified with primer pairs P15/P34 using pUC19/HP1021_C51A-C56A as a template. The PCR fragment was digested with BamHI and XhoI and cloned into pET28Strep using the same enzymes, yielding pET28StrepCemRC51A-C56A.

**Construction of pori2_1 plasmid containing *H. pylori oriC2* region**

To construct the pori2_1 plasmid, the *oriC2* region was amplified by PCR using P32-P33 primer pairs and an *H. pylori* 26695 chromosomal DNA template. The PCR fragment was digested with EcoRI and PstI and cloned into pOC170 using the same enzymes, yielding pori2_1.

### Size-exclusion chromatography with multi-angle light scattering (SEC-MALS)

SEC-MALS was performed using an AKTA Purifier (Amersham) chromatography system equipped with a Superdex 200 Increase 10/300 column (GE Healthcare) and connected to in-line detectors: a MALS detector, DAWN 8+ (Wyatt), and a differential refractometer, Optilab T-rEX (Wyatt). UV absorbance was monitored at 280 nm. Samples were run at room temperature at a flow rate of 0.75 ml/min in Tris buffer (100 mM Tris HCl pH 8.0, 300 mM NaCl), supplemented with 10 mM TCEP when necessary. The results were analyzed using Astra 6 software (Wyatt) in accordance with the manufacturer’s instructions.

**Supplementary Tables**

**Table S1. Strains, plasmids, and proteins used in this study.**

| Strain | Relevant features | Reference/source |
|---|---|---|
| *E. coli* DH5α | *supE44*, *hsdR17*, *recA1*, *endA1*, *gyrA1*, *gyrA96*, *thi-1*, *relA1* | [5] |
| *E. coli* BL21 | F-, *ompT*, *hsdS (rB-, mB-)*, *gal*, *dcm* | GE Healthcare |
| *H. pylori* 26695 | Parental strain | [6] |
| *H. pylori* N6 | Parental strain | [7] |
| *H. pylori* N6 ΔHP1021 | Δ*HP1021*::*aphA-3*; N6 with *HP1021* exchanged to *aphA-3* cassette | [3] |
| *H. pylori* N6 COM/HP1021 | (Δ*HP1021*::*aphA-3*):Δ*HP1021-cat*); N6 Δ*HP1021* in which *aphA-3* was exchanged to *HP1021* and *cat* cassette | [3] |
| *H. pylori* N6 C51A-C56A | Δ*HP1021*::*HP1021*C51A-C56A-*aphA-3*; N6 with *HP1021* exchanged to mutated HP1021 producing C51A-C56A *HP1021* protein variant | This work |
| *H. pylori* N6 C27A | Δ*HP1021*::*HP1021*C27A-*aphA-3*; N6 with *HP1021* exchanged to mutated HP1021 producing C51A-C56A *HP1021* protein variant | This work |
| N6 ΔCys | Δ*HP1021*::*HP1021*ΔCys-*aphA-3*; N6 with *HP1021* exchanged to mutated HP1021 producing cysteine-less *HP1021* protein variant | [3] |
| *C. jejuni* 11168 | *C. jejuni* wild-type strain | [8,9] |

| Plasmid | Relevant features | Reference/source |
|---|---|---|
| pUC19 | Cloning vector, $Amp^R$ | Thermo Fisher Scientific |
| pET28Strep | A pET28a(+) derivative, used for expression of recombinant proteins with an N-terminal Strep-tag II (WSHPQFEK), $Kan^R$ | [3] |
| pGEX-6P-2 | Expression vector for the synthesis of the recombinant proteins with a GST-tag at the N-terminus, $Amp^R$ | GE Healthcare |
| pTZ57R/TΔHP1021 | pTZ57R/T derivative containing HP1021 flanking regions for allelic exchange of *HP1021* for *aphA-3* | [10] |
| pUC18HP1021com | pUC18 derivative containing *HP1021* flanking regions and HP1021 for allelic exchange of *aphA-3* for HP1021-*cat* | [3] |
| pTZ57R/TΔHP1021_1 | pTZ57R/TΔHP1021 derivative containing HP1021 flanking regions and BglII/XhoI sites for cloning of HP1021 mutated genes and subsequent allelic exchange of HP1021 for HP1021 mutated variants | This work |

| | | |
|---|---|---|
| pTZ57R/THP1021C27A | pTZ57R/TΔHP1021_1 derivative vector containg HP1021 C27A mutant alle (TGC→GCT) for exchange of HP1021 wild-type gene for HP1021 C27A variant *via* homologous recombination | This work |
| pTZ57R/THP1021C51A-C56A | pTZ57R/TΔHP1021_1 derivative vector containg HP1021 double mutation C51A and C56A mutant alle (each TGC→GCC) for exchange of HP1021 wild-type gene for HP1021 C51A-C56A variant *via* homologous recombination | This work |
| pET28StrepHP1021 | pET28Strep derivative containing the HP1021 gene, used for the synthesis of CemR wild-type protein | [3] |
| pET28StrepHP1021 NΔCys | pET28Strep derivative containing the HP1021 gene in which codons for cysteine residues C27, C51, and C56 were mutated for alanine codons (TGC→GCT, TGC→GCC ,TGC→GCC, respectively) used for the synthesis of CemR RD-CA variant | [3] |
| pET28StrepHP1021 CΔCys | pET28Strep derivative containing the HP1021 gene in which codons for cysteine residues C216, C238 and C270 were mutated for alanine codons (TGC→GCA, TGC→GCC ,TGC→GCG, respectively); used for the synthesis of CemR CD/DBD-CA variant | [3] |
| pET28StrepHP1021ΔCys | pET28Strep derivative containing the HP1021 gene in which all codons for cysteine residues (C27, C51, C56, C216, C238, and C270) were mutated to alanine codons (TGC→GCT, TGC→GCC, TGC→GCC, TGC→GCA, TGC→GCC, TGC→GCG, respectively), used for the synthesis of CemR $C_{less}$ variant | [3] |
| pET28StrepCemR$_{RD}$ | pET28Strep derivative containing the 1-384 bp of the HP1021 gene, used for the synthesis of CemR$_{RD}$ variant (1-127 aa) | This work |
| pET28StrepCemR$_{CD/DBD}$ | pET28Strep derivative containing the 355-897 bp of the HP1021 gene, used for the synthesis of CemR$_{CD/DBD}$ variant (119-298 aa) | This work |
| pGexCemR$_{CD/DBD}$ | pGEX-6P-2 derivative containing the 355-897 bp of the HP1021 gene, used for the synthesis of CemR$_{CD/DBD}$ variant (119-298 aa) | This work |
| pET28StrepCemRC27A | pET28Strep derivative containing the HP1021 gene in which the codon for cysteine residue C27 (TGC) was mutated to the alanine codon (GCT), used for the synthesis of CemR C27A variant | This work |
| pET28StrepCemRC51A-C56A | pET28Strep derivative containing the HP1021 gene in which the codons for cysteine residues C51 (TGC) and C56 (TGC) were exchanged for the alanine codon were mutated for the alanine codon (GCC), used for the synthesis of CemR C51A-C56A variant | This work |
| pOC170 | Plasmid carrying the *E. coli oriC* sequence | [11] |
| pori2_1 | A pOC170 derivative containing *H. pylori oriC2* | This work |

| Recombinant proteins | Relevant features | Reference/source |
|---|---|---|
| CemR WT | *H. pylori* wild-type CemR protein, Strep-tagged at N-terminus, purified from *E.* coli, Mw 36.8 kDa | [3] |
| CemR $C_{less}$ | *H. pylori* CemR protein variant in which all cysteine residues (C27, C51, C56, C216, C238, and C270) were substituted with alanine residues, Strep-tagged at N-terminus, purified from *E. coli,* Mw 36.6 kDa | [3] |
| CemR RD-CA | *H. pylori* CemR protein variant in which cysteine residues C27, C51, and C56 were substituted with alanine residues, Strep-tagged at N-terminus, purified from *E. coli*, Mw 36.7 kDa | [3] |
| CemR CD/DBD-CA | *H. pylori* CemR protein variant in which cysteine residues C216, C238, and C270 were substituted with alanine residues, Strep-tagged at N-terminus, purified from *E. coli,* Mw 36.7 kDa | [3] |
| CemR C27A | *H. pylori* CemR protein variant in which cysteine residue C27 was substituted with alanine residue, Strep-tagged at N-terminus, purified from *E. coli*, Mw 36.7 kDa | This work |
| CemR C51A-C56A | *H. pylori* CemR protein variant in which cysteine residues C51 and C57 were substituted with alanine residues, Strep-tagged at N-terminus, purified from *E. coli*, Mw 36.7 kDa | This work |
| $CemR_{RD}$ | *H. pylori* CemR truncated protein variant encompassing 1-127 N-terminal amino acids, Strep-tagged at N-terminus, purified from *E. coli*, Mw 16.4 kDa | This work |
| $CemR_{CD/DBD}$ | *H. pylori* CemR truncated protein variant encompassing 119-298 C-terminal amino acids, Strep-tagged at N-terminus, purified from *E. coli*, Mw 23.1 kDa | This work |
| GST-$CemR_{CD/DBD}$ | *H. pylori* CemR truncated protein variant encompassing 119-298 C-terminal amino acids, GST-tagged at N-terminus, purified from *E. coli,* Mw 48.4 kDa | This work |

**Table S2. Primers used in this study**

| Oligo name | Sequence (5' → 3') |
|---|---|
| P1 | AAGCACACAATCCCGCTCGC |
| P2 | CGGCTACCCTTTTGGCGTGG |
| P3 | ATAAAACCGGCGATCCTTCT |
| P4 | GCTTGGATACCCACATGCTT |
| P5 | CTAGCGGATTCTCTCAATGTCAA |
| P6 | GGAGTACGGTCGCAAGATTAAA |
| P7 | FAM-GGAGTAAGAATAGCTTCGAAT |
| P8 | CATCGATAGGATATCCTGGG |
| P9 | GGAGTAAGAATAGCTTCGAATCTCTATTTTGAAAACCCCTATTT |
| P10 | CTATATTTTTTCAATGGTTTAGTGC |
| P11 | Cy5-GGAGTAAGAATAGCTTCGAAT |
| P12 | Bio-CATGTTTGACAGCTTATCATCG |
| P13 | GCATAAGATCTAGGAGTAAGA |
| P15 | GCGGATCCATGAAAATCTTAATCATTGAAGA |
| P16 | GCCTCGAGTTAAAAGTCTAAATAATTTTCATAAAATCC |
| P17 | GCGGATCCGGATTTTATGAAAATTATTTAGACTTTA |
| P18 | GCCTCGAGTTATTTGCGCGGTAAGTTATATT |
| P19 | GATTTAGGGCATTTTGCTGAGATCATCTCTAGC |
| P20 | GCTAGAGATGATCTCAGCAAAATGCCCTAAATC |
| P21 | GTTTCTTCTAAAGTGGCCACTCAAGGGCGCGCCGAACATTTTGTGCG |
| P22 | CGCACAAAATGTTCGGCGCGCCCTTGAGTGGCCACTTTAGAAGAAAC |
| P23 | GCTCACCACGAGCGGCGATTTG |
| P24 | GCTTTAAAATCTTCATCAAATTGG |
| P25 | P-CCGCTCGAGGGATCCTGACTAACTAGGAGG |
| P26 | P-AGATCTCCTCCTTATCTTTAAAGTTTTTGTTTATAACGC |
| P27 | GTCGACTCTAGAGGATCCCCATTAAAGGGGGGGATATTGG |
| P28 | TGAATTCGAGCTCGGTACCCGTAAAAGAGTTCTTGAATGCAT |
| P29 | GGAGTAAGAATAGCTTCGAATGAATAGCTGCTCTTGAAGCAGC |
| P30 | CGCTATCAAATACCATGCGAC |
| P31 | Bio-GGAGTAAGAATAGCTTCGAAT |
| P32 | GCGAATTCGTCTTAAGCCTTAGAGAAGA |
| P33 | GCCTGCAGGGTTTAGTAAAAGTCATAAATA |
| P34 | GCCTCGAGTTATTTGCGCGGCAAATTATACT |

*FAM, 6-fluorescein amidite; Cy5, Cyanine 5; Bio, biotin; P, phosphate

**Table S3.** Dissociation constants of zincon-analyzed $Zn^{2+}$ complexes with CemR variants at various oxidation states for the $Zn^{2+}$-CemR stoichiometry model (reduced, $K_{dred}$, and oxidized $K_{dox}$) and the $Zn^{2+}{}_2$-CemR model (reduced, $K_{d12red}$, and oxidized, $K_{d12ox}$)[a]. Errors are presented in brackets. Values for air-oxidized (nonreduced) and reduced proteins are given in red and blue, respectively. NA, not applicable because the CD/DBD domain or one domain's cysteine residues are missing.

| CemR variant | $K_d$ [M] | -log$K_d$ | $K_{d12}$ [$M^2$] | -log $K_{d12}$ |
|---|---|---|---|---|
| WT | $1.99\times10^{-8}$ (0.42) | 7.71 (0.09) | $1.84\times10^{-15}$ (1.03) | 14.83 (0.25) |
| WT | $5.30\times10^{-8}$ (0.70) | 7.28 (0.06) | $7.53\times10^{-15}$ (2.61) | 14.16 (0.15) |
| CD/CBD-CA | $6.07\times10^{-8}$ (1.15) | 7.23 (0.08) | NA | NA |
| CD/CBD-CA | $1.75\cdot10^{-7}$ (0.13) | 6.76 (0.03) | NA | NA |
| RD-CA | $1.20\times10^{-7}$ (0.04) | 6.92 (0.02) | NA | NA |
| RD-CA | $1.20\times10^{-7}$ (0.04) | 6.92 (0.01) | NA | NA |
| $CemR_{RD}$ | $1.05\times10^{-7}$ (0.12) | 6.98 (0.05) | NA | NA |
| $CemR_{RD}$ | $4.73\times10^{-7}$ (0.23) | 6.32 (0.02) | NA | NA |
| C27A | $3.99\times10^{-8}$ (0.53) | 7.40 (0.06) | $7.58\times10^{-15}$ (0.51) | 14.12 (0.03) |
| C27A | $8.88\times10^{-8}$ (1.03) | 7.05 (0.05) | $2.02\times10^{-14}$ (0.15) | 13.69 (0.03) |
| C51A-C56A | $1.07\times10^{-7}$ (0.17) | 6.98 (0.07) | $2.52\times10^{-14}$ (0.33) | 13.61 (0.06) |
| C51A-C56A | $1.26\times10^{-7}$ (0.17) | 6.91 (0.06) | $3.06\times10^{-14}$ (0.33) | 13.51 (0.05) |

[a] Dissociation constants are defined as following: $Zn^{2+}$-CemR ⇌ *$Zn^{2+}$ + CemR* ($K_d$ = ($[Zn^{2+}]\cdot$[CemR])/[$Zn^{2+}$-CemR]); $Zn^{2+}{}_2$-CemR ⇌ *2$Zn^{2+}$ + CemR* ($K_{d12}$ = ($[Zn^{2+}]^2\cdot$[CemR])/[$Zn_2{}^{2+}$-CemR]).

**A**

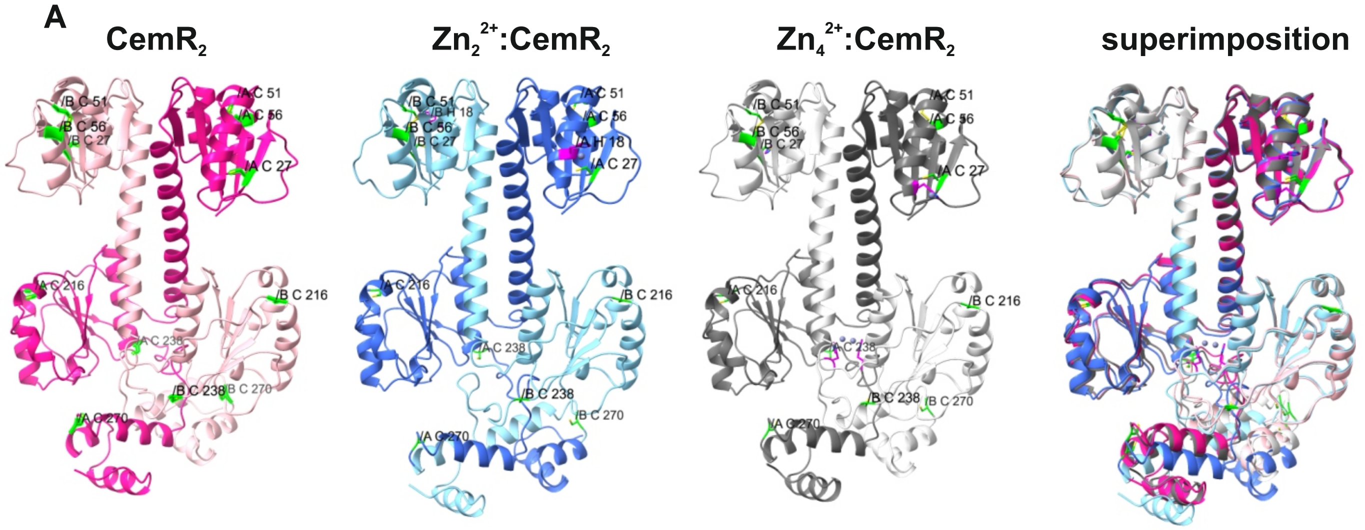

**B**

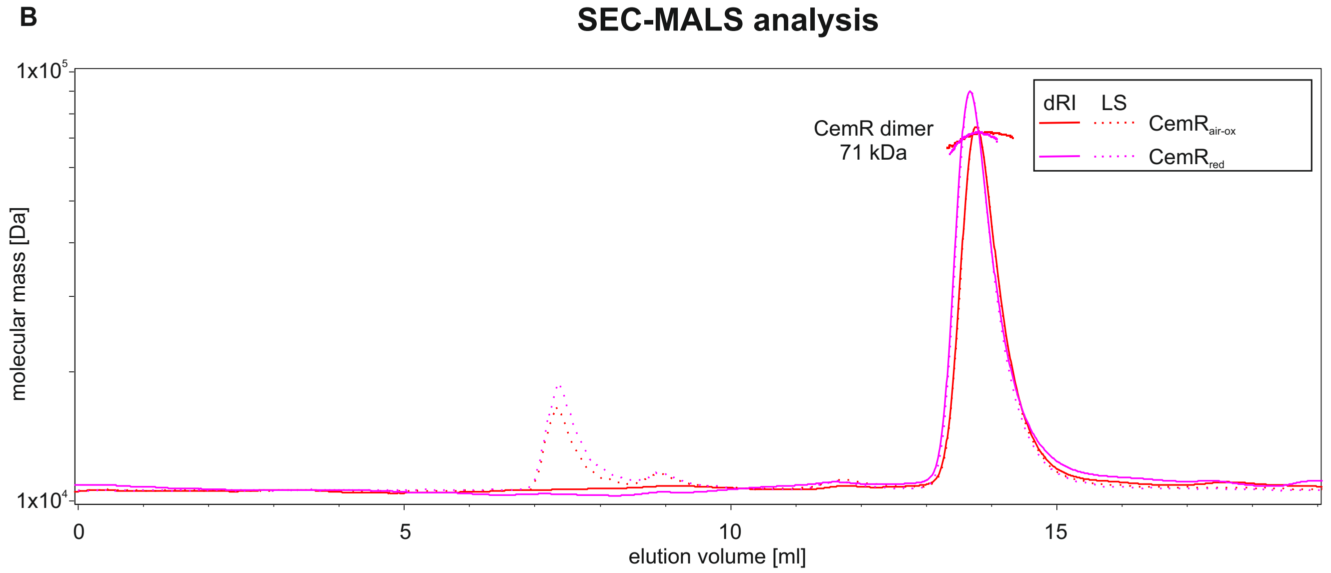

**Fig. S1.** CemR structure modelled by AlphaFold3. (A) A CemR dimer (Uniprot O25665 ) without $Zn^{2+}$ (left hand side, light/dark pink), with $Zn^{2+}$ in 1:1 stoichiometry (middle left, light/dark blue) and 2:1 $Zn^{2+}$:CemR stoichiometry (middle right, light/dark grey) and a superimposition of a CemR dimer with and without $Zn^{2+}$(right hand side) (B) CemR quaternary structure. Unreduced or TCEP-reduced CemR ($CemR_{air\text{-}ox}$ or $CemR_{red}$, respectively) was analysed by SEC-MALS. Only dimers of CemR were observed regardless of redox conditions. LS, light scattering; dRI, differential refractive index.

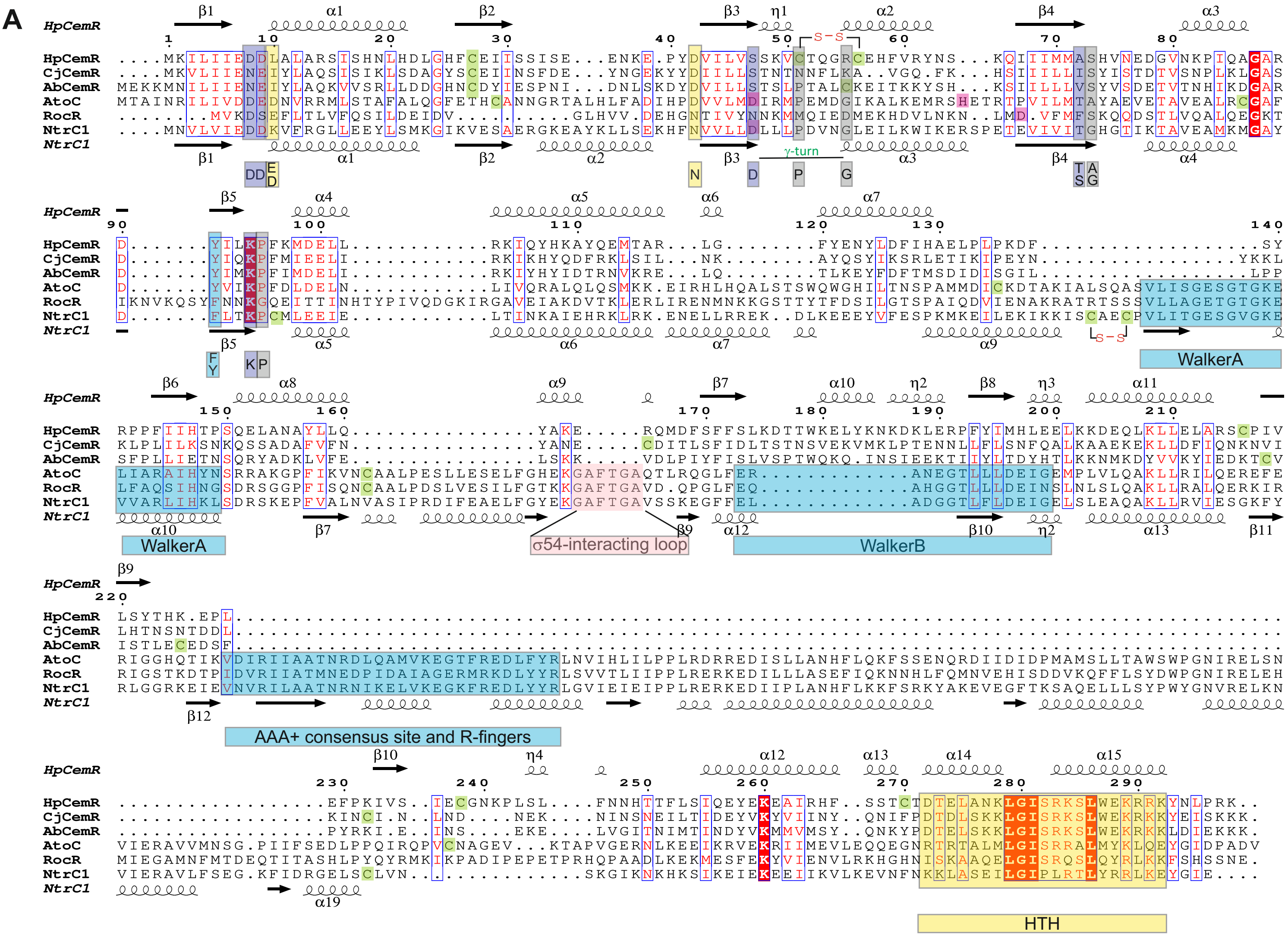

**Fig. S2.** Sequence and structure of HpCemR. Comparison of the HpCemR sequence and its *Campylobacterial* homologues to the most similar RocR and AtoC sequences, and a model representative of NtrC1-type response regulator protein. The sequences of HpCemR (O25665), CjCemR (Q0P820), AbCemR (A8ER38), AtoC (Q06065), RocR (P38022), and NtrC1 (O67198) were used for comparison. Secondary structure predicted in AlphaFold3 for the HpCemR monomer (AF-O25665-F1, presented above HpCemR), or derived from PDB 4L4U structure for NtrC1(AF-O67198-F1, presented below NtrC1). The sequence alignment was performed using Clustal Omega (Madeira *et al.*, doi: 10.1093/nar/gkae241 ) and visualised by ESPript3 (Gouet *et al.*, doi: 10.1093/nar/gkg556). Conservative regions of NtrC-like proteins were marked below the sequence alignment according to Bush *et al.* (doi: 10.1128/MMBR.00006-12); conservative amino acids in the REC domain were marked according to Bourret *et al*. (doi:10.1128/jb.00261-25) and Bourret *et al.* (doi: 10.1016/j.mib.2010.01.015) with the colour coding as in Fig. S3; the HTH domain was marked based on structure prediction. S-S marks disulfide bonds in HpCemR (this work) and NtrC1 (Vidangos *et al*., doi: 10.1002/bip.22333); cysteine residues and phosphorylated amino acids (Grigoroudis *et al.*, for AtoC (doi:10.1016/j.bbagen.2007.04.004), according to Uniprot in RocR and NtrC1) are shaded green and pink, respectively.

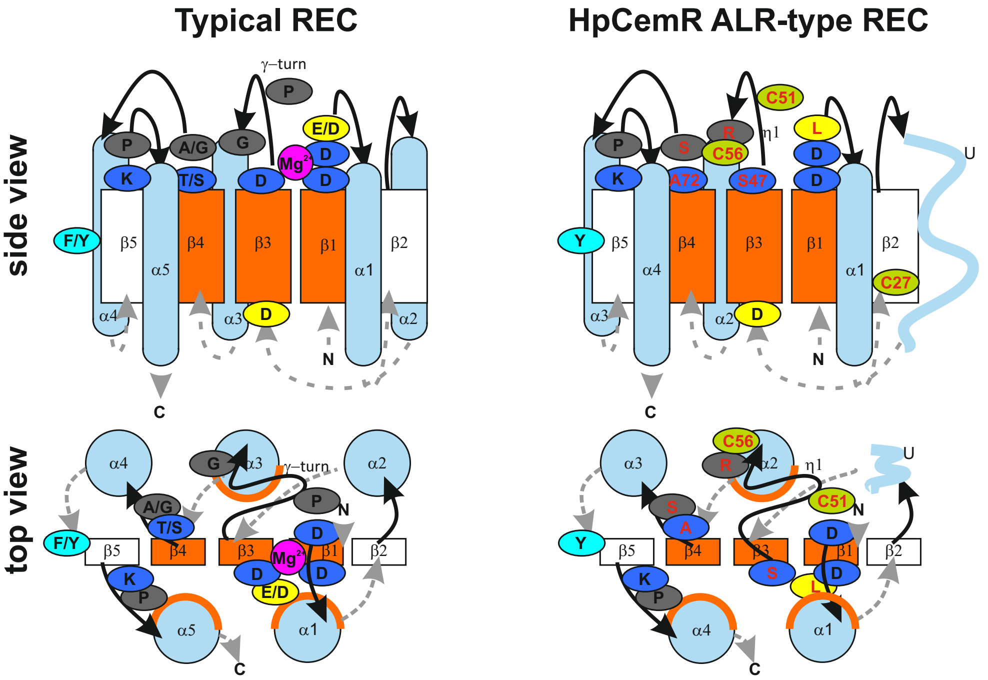

**Fig. S3.** Schematic diagrams of the typical REC and HpCemR REC domains secondary structures. The general structure and amino acids essential for the structure and function of a typical REC were marked as in Bourret *et al.* (doi: 10.1016/j.mib.2010.01.015), and their conservation within the REC domain of HpCemR was compared. Loops connecting β-strands and α-helices are shown as black solid lines on the active site side of the domain and grey dashed lines on the opposite side; arrowheads indicate N-terminal to C-terminal ends. Orange indicates the conserved hydrophobic residues on the central β-strands and faces of three α-helices. The highly conserved residues of the active-site are shown in dark blue, with the moderately conserved aromatic residue in light blue. $Me^{2+}$ (typically $Mg^{2+}$, but not $Zn^{2+}$) is magenta. Residues presumed to be strongly conserved for structural reasons are in grey. Frequently conserved acidic residues of unknown function are in yellow. HpCemR amino acids that differ from those typically found in the REC domain are labelled red. Cysteine residues of the HpCemR REC domain are green. U, unstructured region of HpCemR corresponding to α2-helix in typical REC domains.

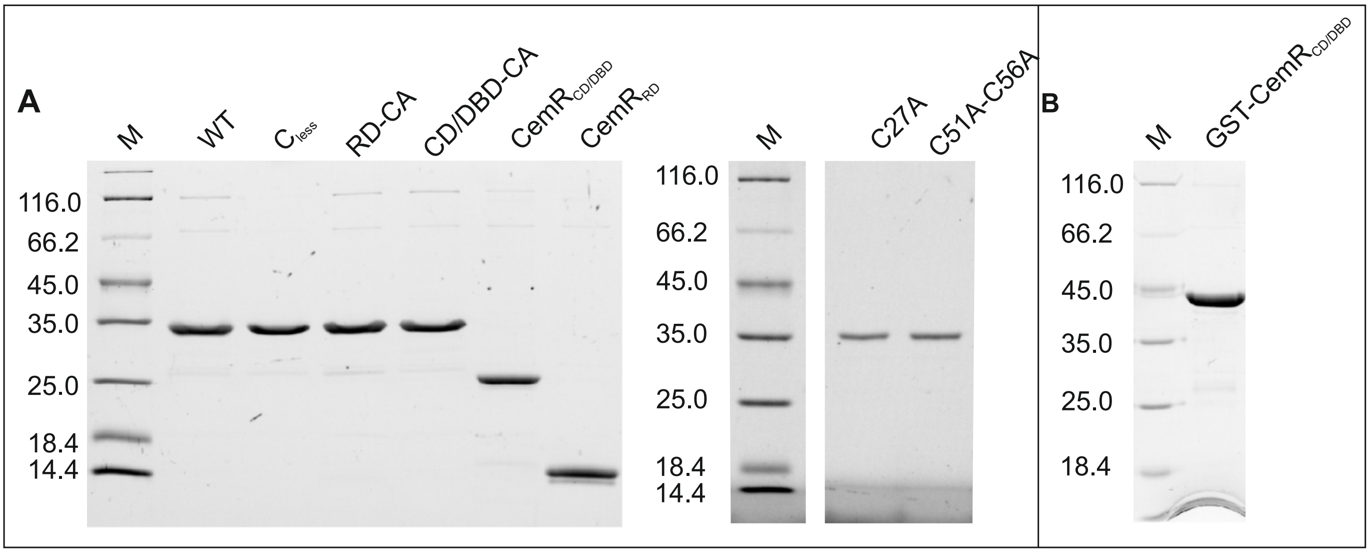

**Fig. S4** (A) The strep-tagged wild-type CemR protein and its variants (Table S1) were purified as described in the Materials and Methods section, according to the same protocol. Each protein variant was separated on a 12% SDS-PAGE gel under reducing conditions and visualised using TCE. (B) The GST-tagged CD/DBD CemR domain (Table S1) was purified as described in the Materials and Methods section, separated on a 10% SDS-PAGE gel under reducing conditions and visualised using TCE. M, molecular weight protein marker (Unstained Protein Molecular Weight Marker, Thermo Scientific, catalog number 26610)

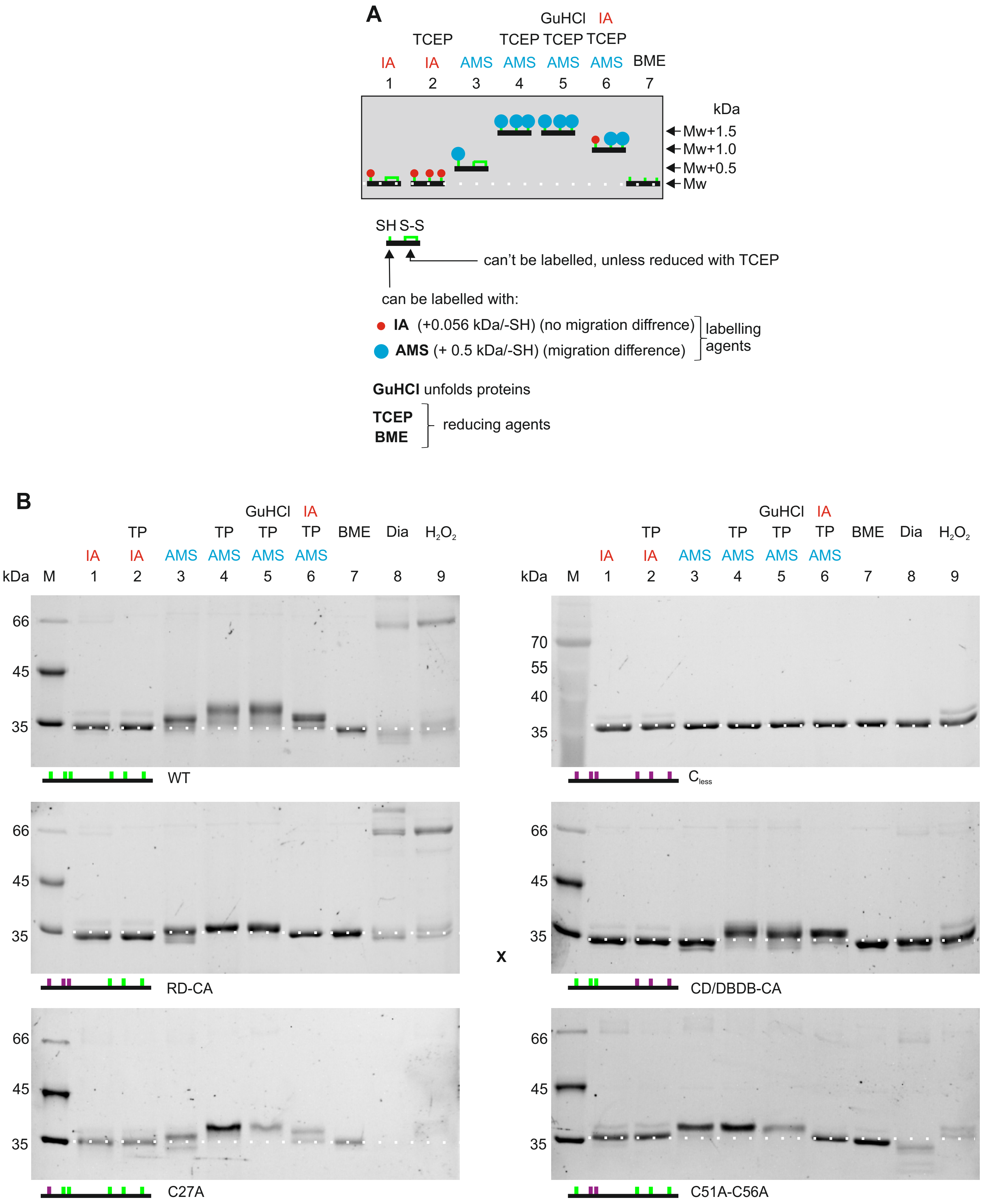

**Fig. S5.** Analysis of the CemR thiol redox state. Schematic overview of the differential thiol-trapping approach (A) and the analysis of the CemR thiol redox state (B). WT and its cysteine variants were incubated in Tris buffer containing reducing and/or labelling agents. GuHCl was added to the denture protein to expose the putative buried cysteine residues to reduction and labelling. The agents were added to the CemR protein sample in subsequent steps, as indicated above the gels. The protein was precipitated with TCA after each step. CemR was resolved in a 10% SDS-PAGE gel under non-reducing conditions and visualized by the TCE method. The white dotted line indicates unlabelled or IA-labelled CemR with no difference in migration. IA, iodoacetamide; AMS, 4-acetamido-4'-maleimidylstilbene-2,2'-disulfonic acid; TP, TCEP; GuHCl, guanidine hydrochloride; BME, beta-mercaptoethanol; Dia, diamide. Digital processing was applied equally across the entire image.

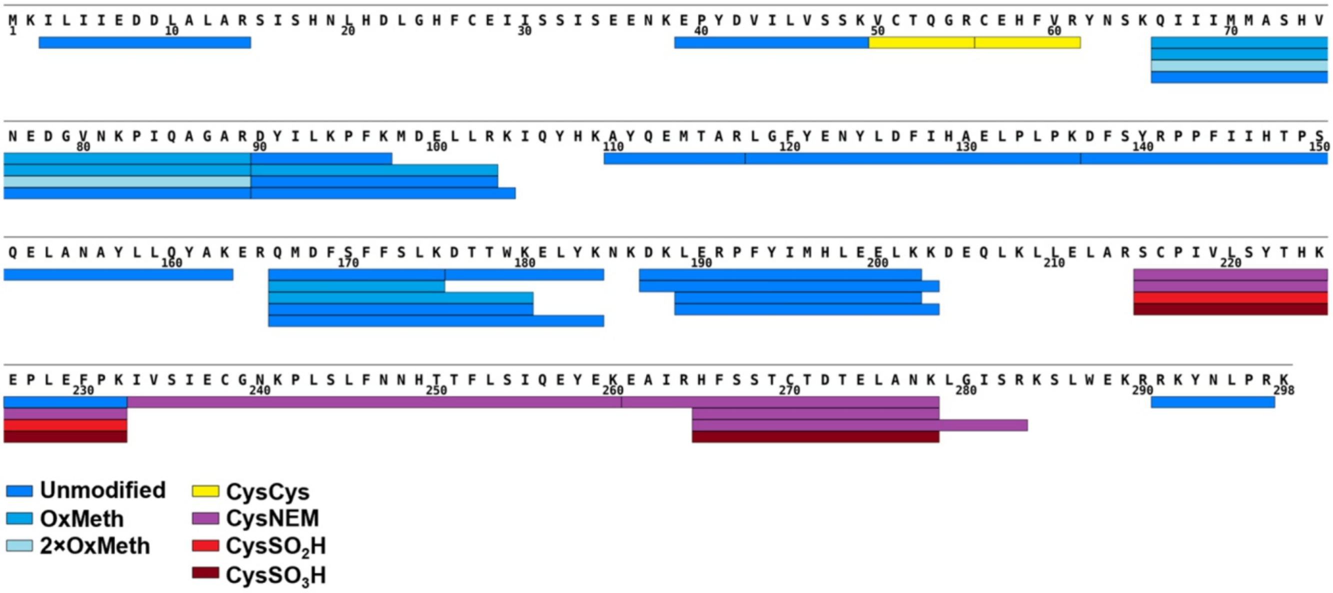

**Fig. S6.** CemR peptide map obtained in bottom-up LC-MS analyses. Cumulatively an 81% CemR sequence coverage was observed. Peptide modification color legend: dark blue — unmodified; blue — oxidized methionine; light blue — 2 oxidized methionines; yellow — disulfide oxidized cysteine; purple — N-ethylmaleimide modified (NEM) cysteine; red — sulfinic acid-oxidized ($SO_2H$) cysteine; dark red — sulfonic acid-oxidized ($SO_3H$) cysteine.

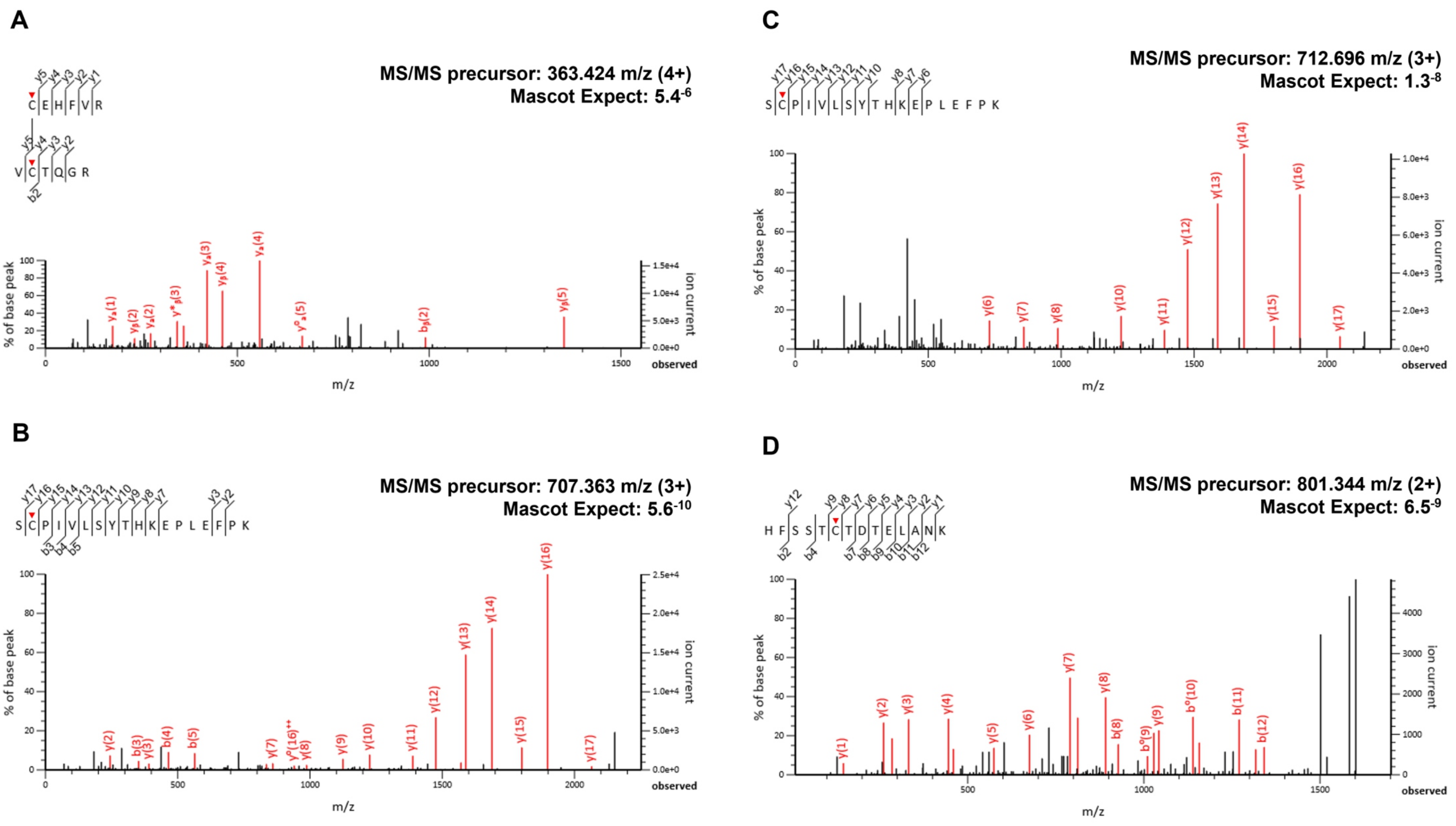

**Fig. S7.** MS/MS spectra of identified CemR peptides containing redox-reactive cysteine residues. (A) C51 to C56 disulfide-linked peptides: VC$_{S-S}$TQGR and C$_{S-S}$EHFVR. (B) C216 sulfinic acid-modified SC$_{SO2H}$PIVLSYTHKEPLEFPK peptide. (C) C216 sulfonic acid-modified SC$_{SO3H}$PIVLSYTHKEPLEFPK peptide. (D) C270 sulfonic acid-modified HFSSTC$_{SO3H}$TDTELANK peptide. Matched b and y fragment ions are indicated in red, unmatched ions are indicated in black. Water neutral losses are symbolized by °. Double-charged fragments are symbolized by $^{++}$. Observed bond cleavages are annotated on sequence in the top left-hand corner of each pane. The m/z and charge of the precursor selected for MS/MS along with Mascot Expect value for the match are indicated in the top right-hand corner of each pane. Cysteine residues bearing the modification are indicated by red pointers.

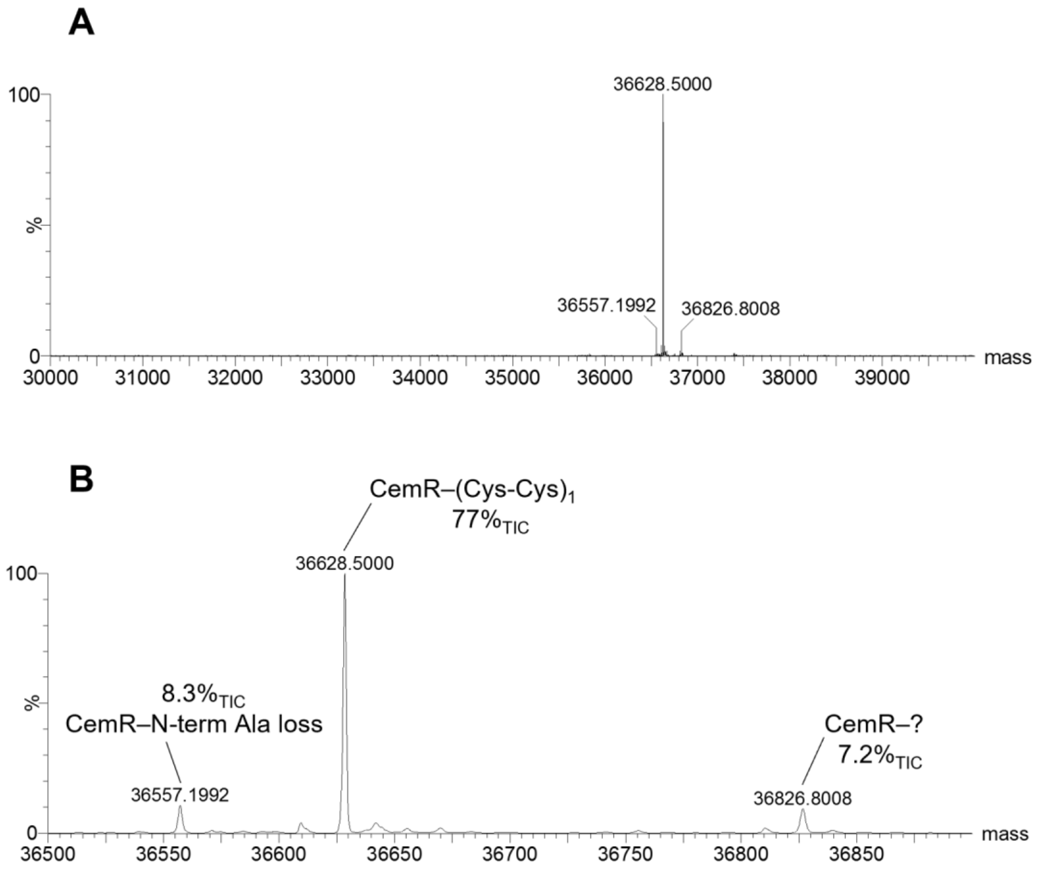

**Fig. S8.** MS spectra of recombinant wild-type CemR after MaxEnt1 deconvolution. (A) Full mass range (30–40kDa). (B) Zoom onto the ion cluster (36.5–35.9kDa). The main peak, which corresponds to CemR mass with a single disulfide, as well as an N-terminal alanine loss in-source fragment, and one unknown species of higher mass are annotated. Peak area as percentage of TIC is also indicated.

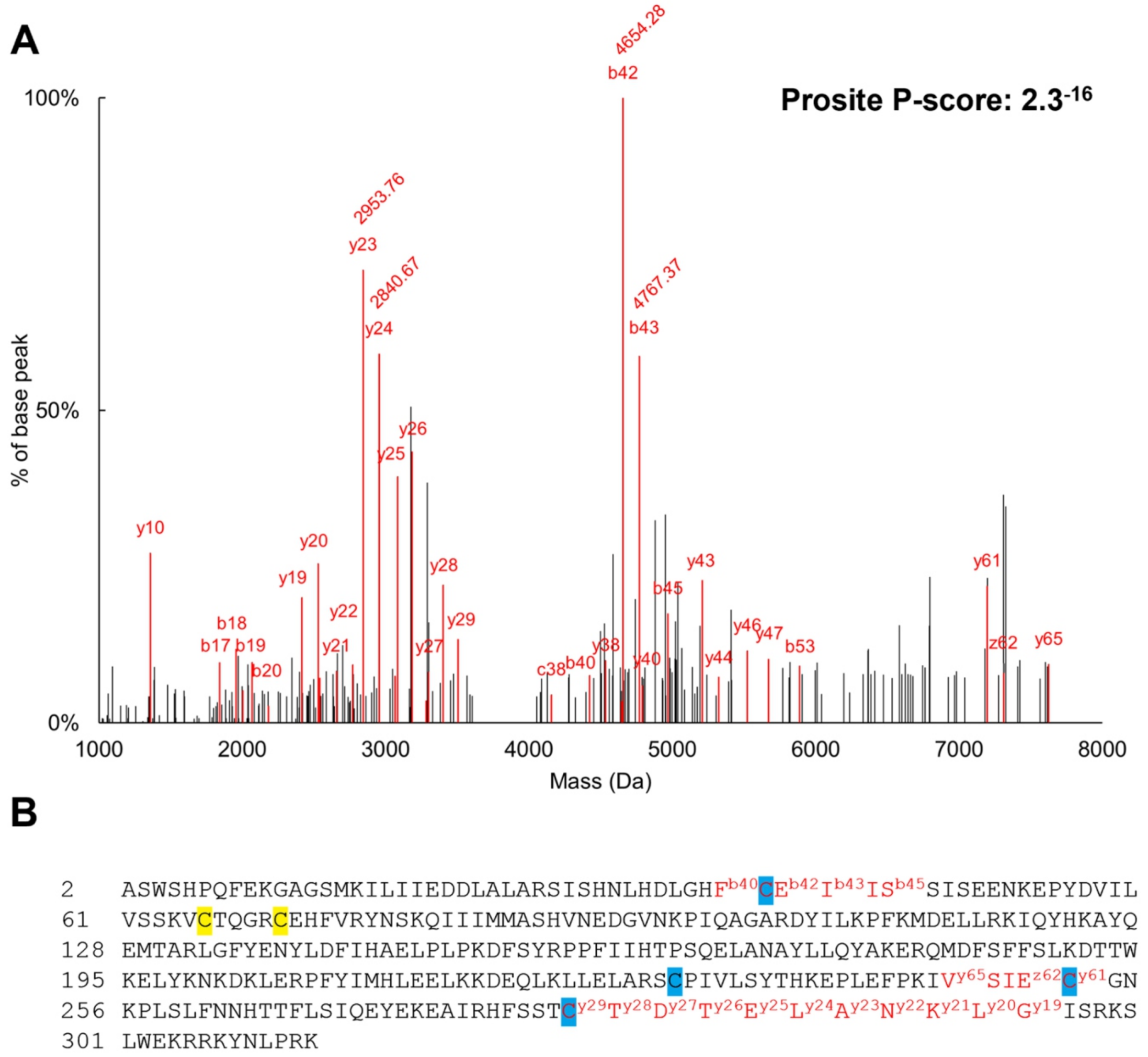

**Fig. S9.** Top-down MS/MS analysis of wild-type CemR. The 44+ charge state ion (833.45 m/z) was CID fragmented with a CE ramp of 25–35V. (A) obtained fragment mass distribution after MaxEnt3 deconvolution and binning of mobility intervals. Peaks matched to CemR sequence BY and CZ ions at ±25 ppm are in red. Unmatched peaks are in black. Ion numbering is according to sequence without the processed N-terminal methionine. Mass of peaks >50% of base peak is annotated. (B) results annotated on CemR sequence. The processed N-terminal methionine residue (pos. 1) is omitted. Cysteine residues were set as Hydrogen Loss (highlighted in yellow) or Unmodified (highlighted in blue) for ion matching. Obtained fragment ion series covering CemR cysteines are in red. Ion numbering is according to sequence without the processed N-terminal methionine.

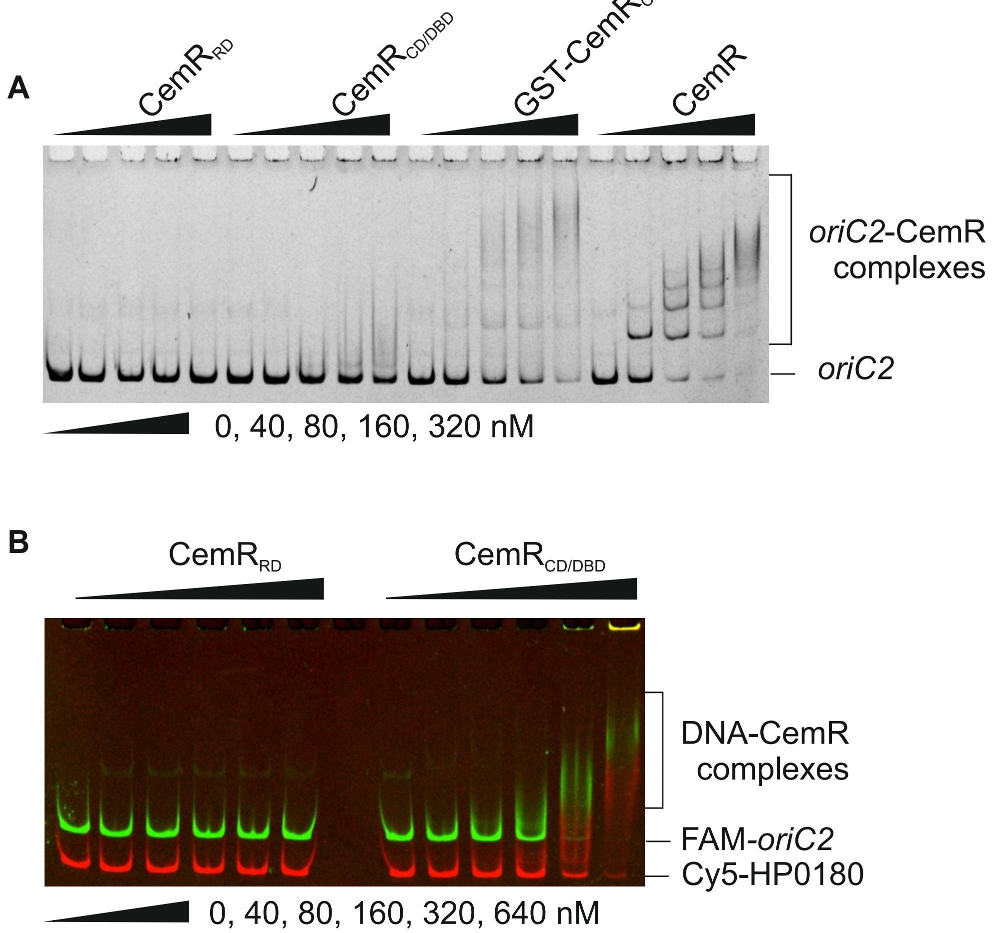

**Fig. S10.** Comparison of the affinity and DNA-binding specificity of CemR variants. EMSA was performed using the specific Cy5-labeled *oriC2* (A); the unspecific FAM-labelled HP0180 gene fragment was added to test the specificity of CemR$_{CD/DBD}$ (B). The CemR$_{RD}$ did not bind DNA, while the CemR$_{CD/DBD}$ affinity and specificity were highly reduced when compared to the CemR wild-type protein (for the specificity of the wild-type CemR see Fig. S9 in Szczepanowski *et al.* (doi: 10.1093/nar/gkab440)).

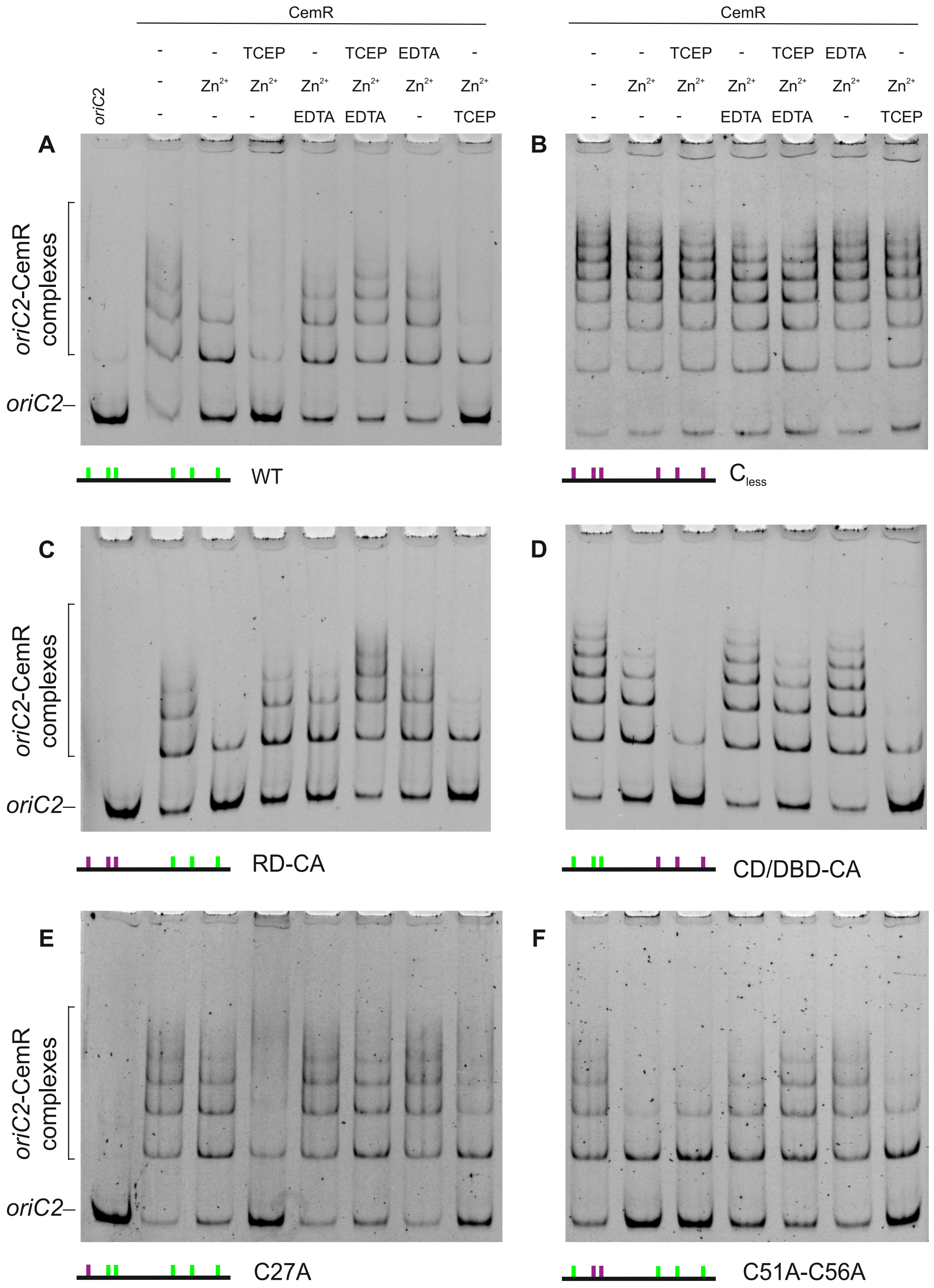

**Fig. S11.** Analysis of the $Zn^{2+}$ influence on CemR-*oriC2* interactions. WT and its cysteine variants (80 nM) were mixed with *oriC2* in Tris buffer and incubated at 37°C (Material and Methods). $Zn^{2+}$ (0.39 μM), TCEP (5 mM), EDTA (1 mM) or $H_2O$ (marked as „-") were added to the CemR-*oriC2* complexes in subsequent steps with 15 min intervals between each supplement, as indicated above the gels (from top to bottom). Complexes were visualised using EMSA. Digital processing was applied equally across the entire image, including controls.
